# The molecular basis of macrochaete diversification highlighted by a single-cell atlas of *Bicyclus anynana* butterfly pupal forewings

**DOI:** 10.1101/2023.08.23.554425

**Authors:** Anupama Prakash, Emilie Dion, Antónia Monteiro

## Abstract

Butterfly wings display a large diversity of cell types, including scale cells of different colors, shapes, and sizes, yet the molecular basis of such diversity is poorly understood. Scales are single-cellular projections homologous to sensory bristles in other arthropods, and different scale types are often found intermingled on the wing, alongside other sensory and epidermal cell types, making the analysis of the complete transcriptomes of each scale type difficult to achieve. To explore scale cell diversity at a transcriptomic level we employed single-cell RNA sequencing of ∼5200 large cells (>6 µm) from 22.5-25-hour male pupal forewings of the butterfly *Bicyclus anynana*. Using single-cell unsupervised, transcriptome clustering, based on shared and differentially expressed genes, followed by *in situ* hybridization, immunofluorescence, and CRISPR/Cas9 editing of candidate cell type specific genes, we annotate various cell types on the wing. We identify the molecular identities of non-innervated scale cells at various stages of differentiation, innervated sensory cell types, and pheromone producing glandular cells. We further show that *senseless*, a zinc finger transcription factor, and *HR38*, a hormone receptor, determine the identity, size, and color of different scale cell types and are important regulators of scale cell differentiation.

**Graphical abstract:** 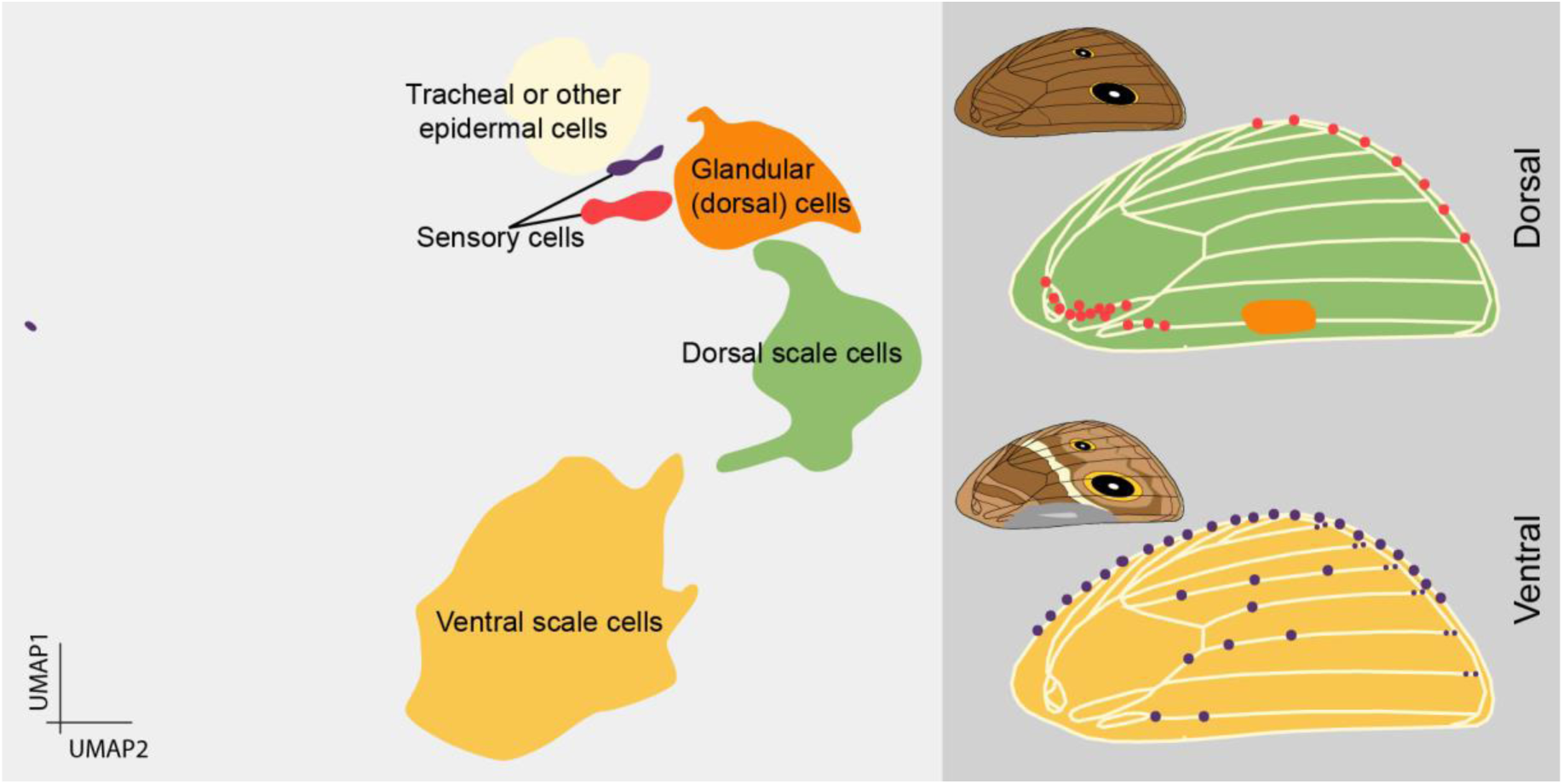

## Introduction

Butterfly wings contain exquisite natural mosaics of individual scale cells that exhibit incredible variation in size, shape, morphology, color, and function. Wing scales are homologous to sensory bristles found in other arthropods (Galant et al., 1998; Zhou et al., 2009) but, in butterflies, these macrochaetes differentiate into paddle-shaped, non-innervated projections, organized in neat rows of alternating larger cover scales overlapping smaller ground scales. Most scales that cover the wing membrane die upon adult emergence leaving behind their colored chitinous skeletons to serve a visual-signal function (Ghiradella, 1991; Greenstein, 1972; Overton, 1966). While scale cell color diversity is the most prominent feature of butterfly wings, wings also display a variety of other cell types that differentiate at multiple locations. Innervated sensory cells, often placed on top of veins (Dickerson et al., 2014; Yoshida and Emoto, 2011; Yoshida et al., 2001), are involved in mechanosensory or chemosensory functions (Aiello et al., 2021; Fabian et al., 2022; Hartenstein and Posakony, 1989) and glandular secretory cells are responsible for producing and secreting sex pheromones, mostly in males (Dion et al., 2016). How such wing cell type diversity is genetically and developmentally regulated remains largely unknown.

Macrochaete developmental progression has been well studied in the fruit fly, *Drosophila melanogaster* (Artavanis-Tsakonas and Simpson, 1991; Furman and Bukharina, 2008; Hartenstein and Posakony, 1990; Lees et al., 1942; Miller et al., 2009; Schweisguth, 2015) (See Supplementary Text) (Fig 1A) and the primary stages of lepidopteran scale development parallel the sensory bristle program, with certain modifications. Scales derive from a single sensory organ precursor (SOP) cell that is specified around 7% of pupal development (PD; 12-15 hours after pupation) (Dinwiddie et al., 2014; Galant et al., 1998). As in flies, SOP cell spacing and organization is determined by a Notch-mediated lateral inhibition mechanism (Reed, 2004). Unlike sensory bristle development, however, after the first cell division, one of the daughter cells (pIIb) dies (at ∼17 hours after pupation), eliminating the neuron and sheath progeny cells (Galant et al., 1998). By ∼14% PD (∼24 hours after pupation), neat rows of scale cell precursors (pIIa cells) are seen. In *J. coenia* butterflies, these pIIa cells express a butterfly homolog of the AS-C genes, *ASH1 (achaete-scute homolog 1)* (Galant et al., 1998), which is essential for scale development (Zhou et al., 2009). Division of the pIIa cell gives rise to a scale and socket cell (Fig S1A) (Dinwiddie et al., 2014). Alternating cover and ground scales can be distinguished based on size by ∼21% PD (Dinwiddie et al., 2014).

**Fig 1:**
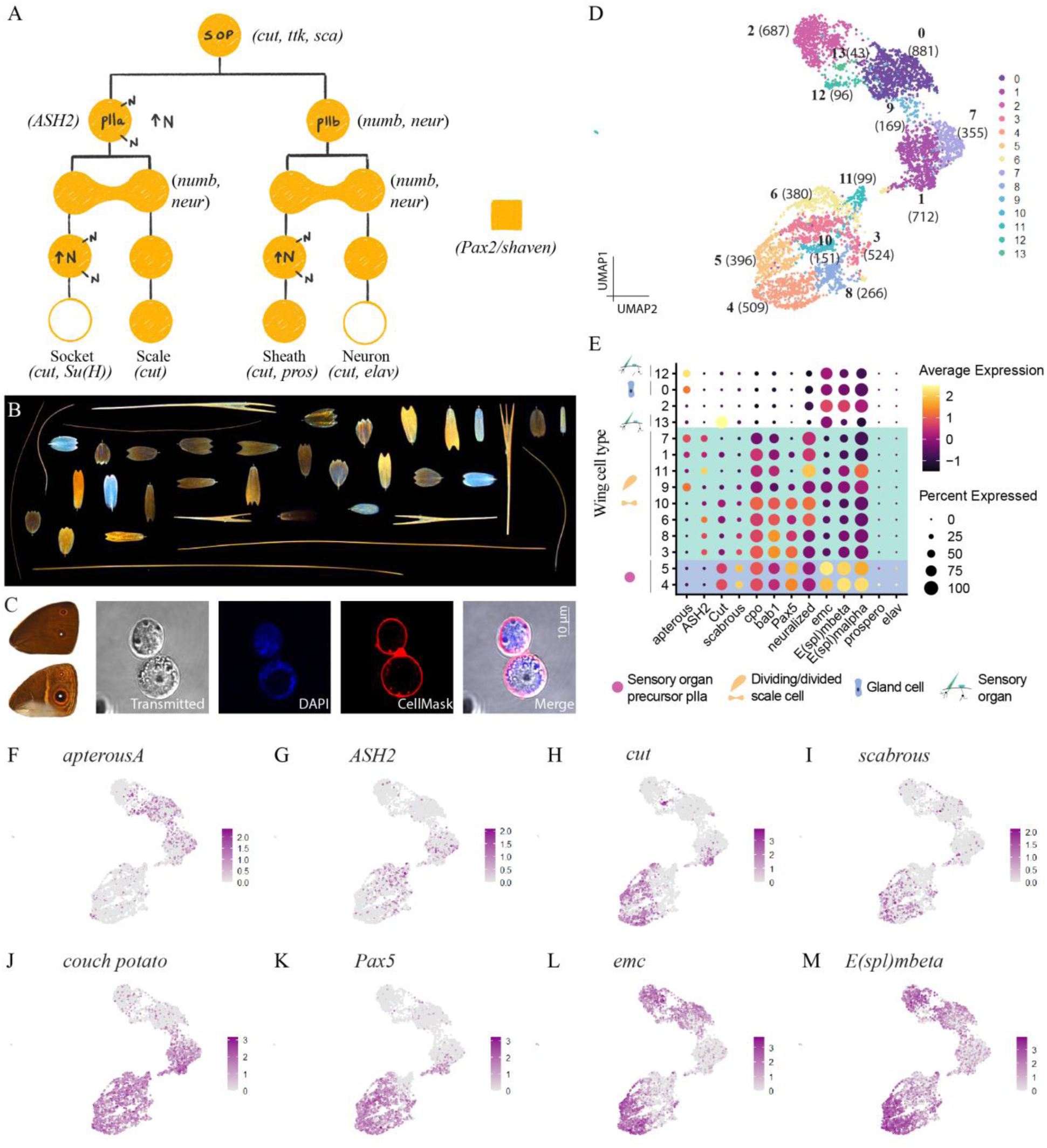
Scale cell diversity and a single-cell atlas of 24-hour male pupal forewings of *Bicyclus anynana*. (A) A schematic of the sensory bristle cell lineage of *Drosophila melanogaster* (see Supplementary text). Genes expressed at different stages along the cell lineage are indicated in parentheses. *D. melanogaster Pax2/shaven* is expressed in the SOP, pIIa and pIIb cells (filled circles) but in the following cell divisions, its expression is restricted to the scale and sheath cells only (open circles). N: Notch receptor and increased Notch signaling. (B) Diversity in scale color, morphology and size seen on the forewings of male *B. anynana* butterflies. (C) Male dorsal (top) and ventral (bottom) forewings of *B. anynana* and isolated single cells from a 24- hour pupal forewing stained with DAPI and CellMask Plasma Membrane stain. (D) A Uniform Manifold Approximation and Projection (UMAP) plot of 5268 large pupal forewing cells separated into 14 distinct clusters. The number of cells in each cluster is shown in brackets. (E) Dotplot of different marker gene expressions within each cluster. Putative higher-order cell classes are indicated by the schematics. The horizontal blue and green boxes highlight the putative scale precursor cells and the dividing/divided scale cells, respectively. (F-M) Expression of various genes used to annotate dorsal vs ventral cells and identify scale and scale cell precursors.

Given the established homology of sensory bristles and butterfly scales, and an understanding of the initial stages of scale development, we sought to explore how scales varying in morphology, color, and size (Fig 1B) vary in their differentiation program. In addition, we aimed to identify and characterize other large wing cell types beyond scale cells. We employed single-cell transcriptomics to collect gene expression information from individual large cells of developing pupal forewings of *Bicyclus anynana* butterflies. We obtained fourteen biologically meaningful cell populations using unsupervised clustering of ∼5200 large cells and annotated the scale cell clusters using known marker genes from the extensive *Drosophila* sensory bristle literature (Supplementary Text). We further annotated epidermal and sensory cell types on the wings such as the marginal mechanosensory bristles and the glandular cells that produce male sex pheromones. By performing functional characterization of highly expressed genes within the broad scale cell clusters, we uncovered important roles for the genes *senseless (sens)* and *hormone receptor 38* (*HR38*) in regulating scale characteristics and identities of scale cell types.

## Results

### A single-cell atlas of 24-hour male pupal forewings of *Bicyclus anynana* and the identification of scale cell clusters

We applied single-cell transcriptomics to 22.5-25-hour male pupal forewings of *B. anynana,* which corresponds roughly to 14-16% of pupal development (PD) (Fig 1C). We dissociated pupal wing tissues to obtain single cells and sorted these cells based on size and viability (Fig 1C) (Prakash and Monteiro, 2020a). We collected cells larger than ∼6 µm in diameter. At ∼24 hours, cells of this size or larger correspond to the pIIa cells, their progeny (Galant et al., 1998), and other large cells on the wing, based on confocal images of wings from that timepoint stained with DAPI and CellMask Plasma Membrane stain (Fig 1C, S1B). Cells were processed via the 10X Genomics microfluidics system (10x Genomics Single cell 3’ mRNAseq V3) to generate the single-cell libraries. After quality control, 5268 high-quality single-cell transcriptomes were processed downstream (see Materials and Methods) to obtain a single-cell atlas consisting of 14 cell clusters that we deemed to provide a biologically meaningful resolution of the large wing cell types (Fig 1D).

Within the wing cell atlas, we could first distinguish between dorsal and ventral cells based on the expression of the dorsal marker gene *apterous A (apA)* (Fig 1E, F) (Prakash and Monteiro, 2018). Around 42% of the cells analyzed i.e., 2,213 cells belonging to clusters 0, 1, 7, 9 and 12 had high levels of expression of *apA* (Fig 1E, F).

We next investigated the molecular identity of the scale cells and their diversity, and then characterized the other cell types on the wing. We deduced that cells in clusters 1 and 3-11 are future scale cells at different time points in their differentiation from a sensory organ precursor (pIIa) to a differentiated scale and socket cell (Fig 1A, D, E). These clusters expressed a neural cell type marker, *couch potato* (*cpo*) (Fig 1J) (see Supplementary text for references from *Drosophila* literature), and *bric a brac 1* (*bab1*) (Fig S2), a gene known to mark both cover and ground scales in butterflies (Ficarrotta et al., 2022). Clusters 4 and 5 are likely pIIa cells at earlier time points along the scale cell lineage, while cells in clusters 1, 3, 6-11 are dividing/divided scale cells. The early pIIa cells expressed neural markers and high levels of *extramacrochaetae (emc)* and *enhancer of split mbeta (E(spl)mbeta)* which are downstream targets of Notch signaling (Fig 1E; horizontal blue box, L, M). In contrast, cell clusters 1, 3, 6-11 showed reduced levels of Notch signaling alongside expression of some neural markers suggesting that they are pIIa cells at later stages of the scale cell lineage or, the progeny of the pIIa cell (Fig 1E, horizontal green box). This is supported by the higher levels of *neur* expression in cells of clusters 1, 6, 7, 10 and 11 (Fig 1E). *neur* increases over time in the pIIa cell and is asymmetrically distributed into a single progeny, the scale cell, in the following cell division. None of the cell types expressed pIIb markers *elav* and *prospero* corresponding with the death of pIIb cells during lepidopteran scale cell development (Fig 1E).

Cells in clusters 1 and 3-11 also expressed genes such as *cut, scabrous (sca), tramtrack (ttk)* and *sanpodo (spdo)* (Fig 1E, H, I, S2). *spdo* had a broad expression across clusters 1 and 3-11, similar to *cpo* and *bab1* (Fig S2), while a large number of cells, predominantly in clusters 4 and 5, expressed high levels of *cut*, *sca, ttk* and a zinc finger transcription factor *basonuclin 2*, related to *Drosophila disco* (Vanhoutteghem et al., 2009) (Fig 1E, H, I, S2)*. cut, sca* and *ttk* expression overlapped extensively with a putatively annotated *Ba-Pax5* gene, which is a homolog of *Drosophila Pax2/shaven* (Fig 1K), a gene essential for bristle development in *Drosophila* (Kavaler et al., 1999). Cells in a subset of the above clusters (1, 3, 6, 7, 8 and 11) also expressed *BaASH2* (Fig 1G), whose homolog has a functional role in scale development of *Bombyx mori* (Zhou et al., 2009) (Fig S3).

### Cell clusters express a combination of various gene expression programs indicating they contain cells at different stages of differentiation

We next sought to identify groups of significantly co-expressed genes (gene expression programs or GEPs), akin to different ‘cellular processes’, and infer their relative contributions to each of the previously identified clusters (Kotliar et al., 2019). A GEP exclusively used in one cell type would indicate an identity GEP, while GEPs expressed in multiple cell types would be indicative of common cellular programs like the cell cycle (Kotliar et al., 2019). We employed consensus non-negative matrix factorization (cNMF) and identified 13 GEPs that were informative for our analysis, along with their relative contributions to each cell cluster (Fig 2A, S4A-E) (Brückner et al., 2021).

**Fig 2:**
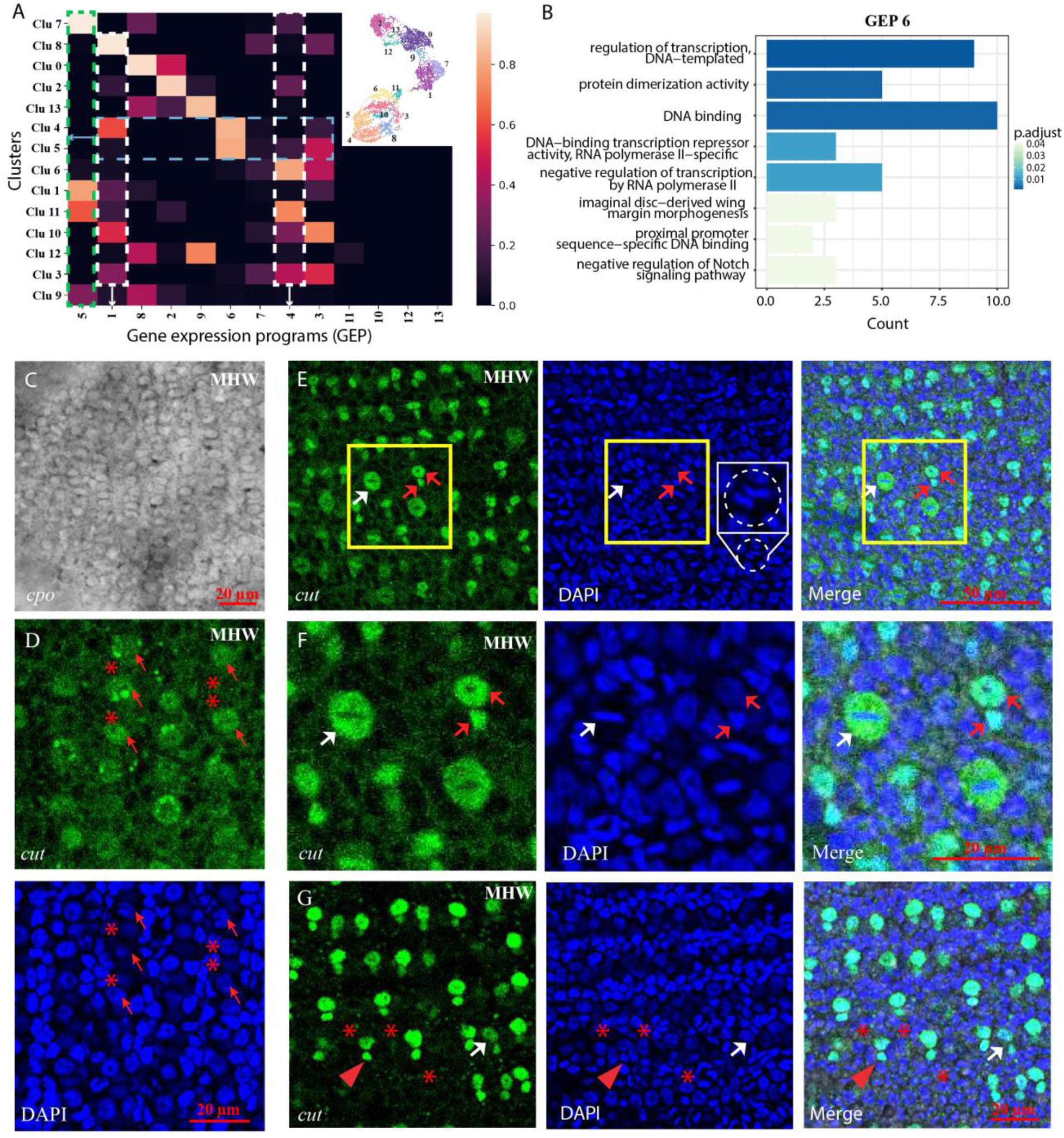
Gene expression programs in cell clusters and expression of *cpo* and Cut. (A) Gene expression program (GEP) usage map indicating the contributions of 13 GEPs to each cell cluster. White and green dashed boxes highlight the usage of GEPs 1, 4 and 5 across different cell clusters. Blue dashed box indicates the usage of multiple GEPs within cell clusters 4 and 5. Inset shows a cropped UMAP plot of the *B. anynana* single-cell atlas with the corresponding cluster numbers marked. (B) Gene enrichment plot of the top 60 genes within GEP 6. (C) *cpo* mRNA expression in ∼25-28-hour pupal wings of *B. anynana*. Expression is seen in cells arranged in neat rows corresponding to the pIIa cells or their progeny. (D-G) Co-immunostaining of ∼24-hour pupal wings of *B. anynana* with DAPI and anti-Cut antibody. (D) Cut protein is seen in the cytoplasm of some pIIa cells (red arrows). Not all pIIa cells express Cut (red stars). (E-F) Cut protein moves from the cytoplasm in pIIa cells to the nucleus in the divided scale and socket cells. Yellow boxed regions are magnified in (F). White arrows indicate Cut protein expression anti-colocalized with nuclear DAPI expression in the pIIa scale precursor cell. Red arrows show nuclear localization of Cut protein expression in the progeny of the pIIa cell, i.e., the scale and socket cells. There is spatial heterogeneity in the scale precursor pIIa cell division across the wing. The white dashed circle in (E), expanded in the inset, highlights a dividing cell. (G) A wing region showing divided scale and socket cell pairs expressing Cut (red arrowheads). Cut protein is however not seen in all scale and socket cells (red stars). White arrow indicates a scale and socket pair expressing lower amounts of Cut protein. MHW: Male hindwing.

Clusters 4 and 5 uniquely expressed GEP 6 (Fig 2A, B), a module enriched in genes involved in regulation of transcription, DNA binding, imaginal disc-derived wing margin morphogenesis and negative regulation of the Notch signaling pathway. GEP 6 consisted of many of the cell fate specification markers discussed above such as *cut*, *ttk* and *Pax5* along with *E(spl), emc* and *basonuclin 2* (Table S1). The exclusive use of GEP 6 in only cells of clusters 4 and 5 suggests that this is an identity GEP, reflecting the early nature of these cells along the scale cell lineage. In combination with GEP 6, clusters 4 and 5 also expressed GEPs 1 and 3, respectively (Fig 2A, blue dotted box). GEP 1 (and GEP 4), widely expressed across scale cell clusters (Fig 2A, white dotted boxes), consisted of numerous ribosomal proteins and ATP synthase subunits suggesting that cells expressing this program were actively growing in either the G1 or G2 phase of the cell cycle (Fig S4F, G). GEP 3, which was also expressed in many cell clusters, was enriched in genes contributing to myosin binding (Fig S4H) as well as cell division (protein MIS12 homolog, protein Spindly) (Table S1), potentially correlating with the mitotic M phase of the cell cycle.

Another identity GEP, GEP 5, was uniquely and highly expressed in clusters 1, 7 and 11 (Fig 2A, green dotted box). GEP 5 comprised of many cuticle proteins, *neur*, a cytoskeleton binding protein, Klarsicht, and a gene related to *Drosophila insensible* that is involved in negatively regulating Notch signaling (Table S1). This GEP also comprised two proteins, Zasp-like and spire, which can bind actin and nucleate new actin filaments respectively (Ashour et al., 2022; Quinlan et al., 2005). This suggested that cells in clusters 1, 7 and 11 are the progeny of pIIa cells, potentially undifferentiated scale cells, rather than socket cells, based on the size criterion used for initial cell sorting. Overall, the combinatorial expression and varying levels of usage of the various GEPs in the different cell clusters indicated that, as predicted, different clusters within the scale cell class were composed of cells at different stages along the scale cell lineage.

We further verified cells of the scale cell lineage (clusters 1, 3-11) by examining the expression of two of the identified marker genes, *cpo* and Cut, in 24-28 hour developing pupal wings of *B. anynana*. *in situ* hybridization of *cpo* mRNA showed restricted expression in the cells of the scale cell lineage (Fig 2C, S5A), and not in the surrounding epidermal cells. Like *cpo*, we observed diffused Cut protein expression in the cytoplasm of pIIa cells (Fig 2D; red arrows). However, Cut was only expressed in some, but not all pIIa cells (Fig 2D; red stars). As the pIIa cells divided into scale and socket cells, Cut protein expression changed from cytoplasmic to nuclear (Fig 2E). Cut cytoplasmic expression, clearly anti-localized with nuclear DAPI (Fig 2E, F; white arrows, S6A), changed to strong nuclear localization in the scale and socket nuclei (Fig 2E, F; red arrows). Dramatic differences in Cut expression patterns correlated with cell cycle progression, which was non-uniform across the wing. Different phases of the cell cycle were evidenced by the shape of the nucleus (circular [red arrow] vs disc-shaped [white arrow] vs separating disc-shaped [dashed circle]) (Fig 2E, F). In regions of the wing where most of the pIIa cells had divided, Cut expression was still non-uniform, being strongly expressed in some scale and socket pairs (Fig 2G; red arrowheads), expressed at lower levels in a few pairs (Fig 2G; white arrow) and not expressed at all in others (Fig 2G; red stars).

### *Hormone Receptor 38* and *senseless* are expressed in the scale cells

We next identified other highly expressed genes in the scale cell clusters, which comprised of the proteins Zasp, Spire, Paramyosin and many non-coding RNAs (Fig S7). We visualized the expression of three highly expressed genes in the dividing/divided scale cells - a *Hormone Receptor 38 (HR38)* (Fig 3A), a zinc-finger transcription factor, *zn271,* orthologous to *Drosophila senseless (sens)* (Fig 3C) and *irregular chiasm C-roughest* (*rst*) (Fig S8). *in situ* hybridization verified the expression of *HR38* and *sens* in neat rows across the pupal wings corresponding to either the scale or scale precursor cells (Fig 3B, D, S5B, C). *rst* was also expressed in the scale cells (Fig S8).

**Fig 3:**
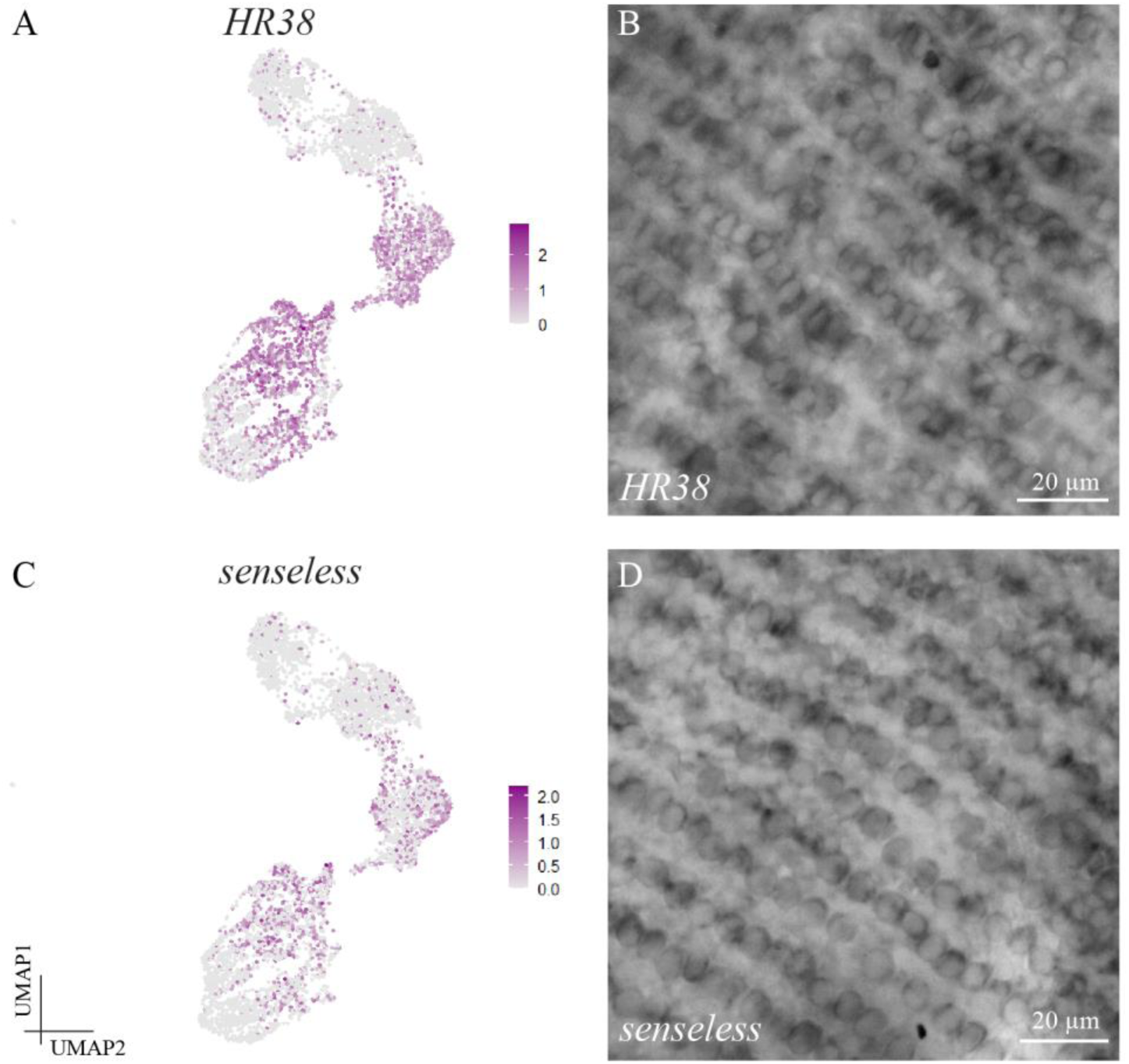
*HR38* and *senseless* are expressed in the scale cells. (A, C) UMAP expression plots of (A) *HR38* and (C) *senseless*. *HR38* (B) and *senseless* (D) mRNA are expressed in neat rows across 24-26 hour pupal wings in the scale building cells.

### *HR38* and *senseless* regulate scale identity, spacing, color and size

We then used CRISPR-Cas9 to functionally verify the role of *BaPax5 (*a homolog of *D. melanogaster Pax2/shaven)* and identify the functions of the highly expressed *paramyosin*, *HR38,* and *sens* genes in scale development. Knockout of *Pax5* in *B. anynana* led to large patches of the wings and body devoid of scales and sockets (Fig 4A, B, S9), as also observed in *Drosophila* bristle development (Kavaler et al., 1999). Similarly, knockout of *paramyosin* led to loss of scales across the wing (Fig S10).

**Fig 4:**
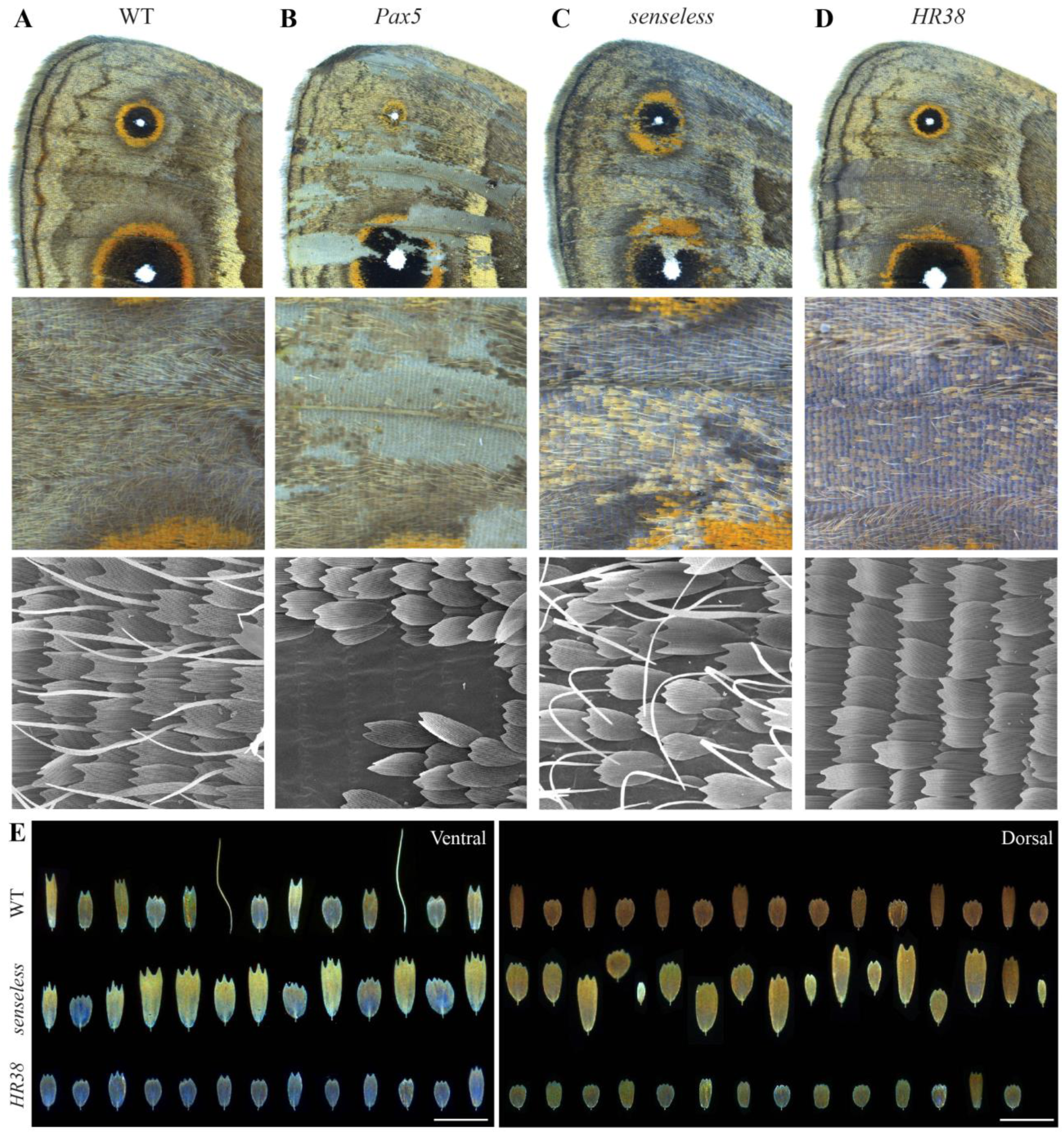
*Pax5, senseless and HR38* are important regulators of scale development. (A) Wildtype (WT) ventral forewing of *B. anynana* at different magnifications. Homologous regions of the wing in (B) *Pax5* (C) *senseless* and (D) *HR38* crispants. Images in row 3 are SEM images. (E) Arrangements of scales along a row in the WT wing compared to scale arrangements in a *senseless* and *HR38* crispant on both the dorsal and ventral surfaces. Lack of a neat, horizontal order in the *senseless* crispants indicates the loss of row arrangement and the disordered nature of the scales. Scale bars: 150 µm.

Knockout of *sens* produced crispants that exhibited changes in scale identity and loss of regular spacing (Fig 4A, C). Within the targeted patches, scales no longer appeared in neat, overlapping rows and were instead non-overlapping, less dense, and randomly arranged (Fig 4C, SEM). Brown, black, beige, or orange ventral cover scales were transformed into large, yellow scales with highly serrated distal edges (Fig 4C, E – ventral, S11) or, in some patches, into rounded silver scales (Fig S11 – Ind 1, red box). Ground scales increased in size (Fig 4E – ventral). On the dorsal surface, there was a similar lower density of scale cells with cover and ground scales increasing in size and becoming yellower (Fig 4E – Dorsal) while some became tiny and silver (Fig S11 – Ind 1 and 9). The tiny scales were disproportionately present on the dorsal wing surfaces compared to the ventral surface. Interestingly, despite the apparent uniform expression of *sens* mRNA across the entire pupal wing, including in the eyespot centers (Fig S5B), the white eyespot center scales appeared to be insensitive to *sens* expression because no transformations of these scales were visible even when surrounding areas were targeted by the knockout (Fig S11 – Ind 3, 8, 9).

Unlike *sens* crispants, knockout of *HR38* led to the loss of all hair-like scales (Fig 4D) indicating that this gene is required to promote hair-like scale identities. In addition, all cover and ground scales on both the dorsal and ventral surfaces decreased in size as compared to wildtype scales (Fig 4E). Cover scales became similar in size to ground scales. Color was also affected, especially in the beige and brown areas of the ventral forewing, that became lighter (Fig 4D, S12). In one crispant, scales were transformed into iridescent silver scales, mostly restricted to the proximal and posterior regions of the wing (Fig S12 – Ind 3). Genotyping results for *Pax5*, *senseless* and *HR38* cripants are shown in Fig S13.

As a result of detailed comparisons of wildtype and cripant wings, we identified an intermediate scale cell type in wildtype wings, amongst the alternating cover and ground scales (Fig S14). This scale cell type was most frequently seen on the ventral surfaces and was additionally found in other species we investigated like *Junonia orithya*, *Junonia almana* and *Catopsilia pomona* (Fig S14). This suggests that scale rows contain an extra cell type beyond ground and cover scales.

### Identification of pheromone producing glandular cells and innervated sensory cell types

Progressing beyond the scale cell clusters, we annotated cells in cluster 0 as the pheromone producing glandular cells of *B. anynana* forewings. These are epidermal cell types based on the high level of Notch signaling in these clusters (Fig 1E, L, M) and the lack of neural markers. GEP 8 was highly expressed in cluster 0 with some expression in clusters 9, 12 and 13 and, consisted of genes involved in metabolic processes, positive regulation of Toll signaling and structural constituent of cuticle (Fig 5A dotted white box, B, Table S1). Some of the highly expressed genes in cluster 0 (and 9, but not 2) also consisted of an odorant binding protein Obp- 56a, various juvenile hormone binding proteins, cuticle proteins, and a trehalose transporter Tret- 1 (Fig 5D, Table S1). Furthermore, a fatty acyl-CoA reductase (FAR), an important enzyme in the sex pheromone biosynthesis pathway (Liénard et al., 2014) was expressed at low levels in a few cells of cluster 0 (Fig S15A). Thus, cells in cluster 0 were active cuticle synthesizing, metabolic cells which were dorsally located based on *apA* expression.

**Fig 5:**
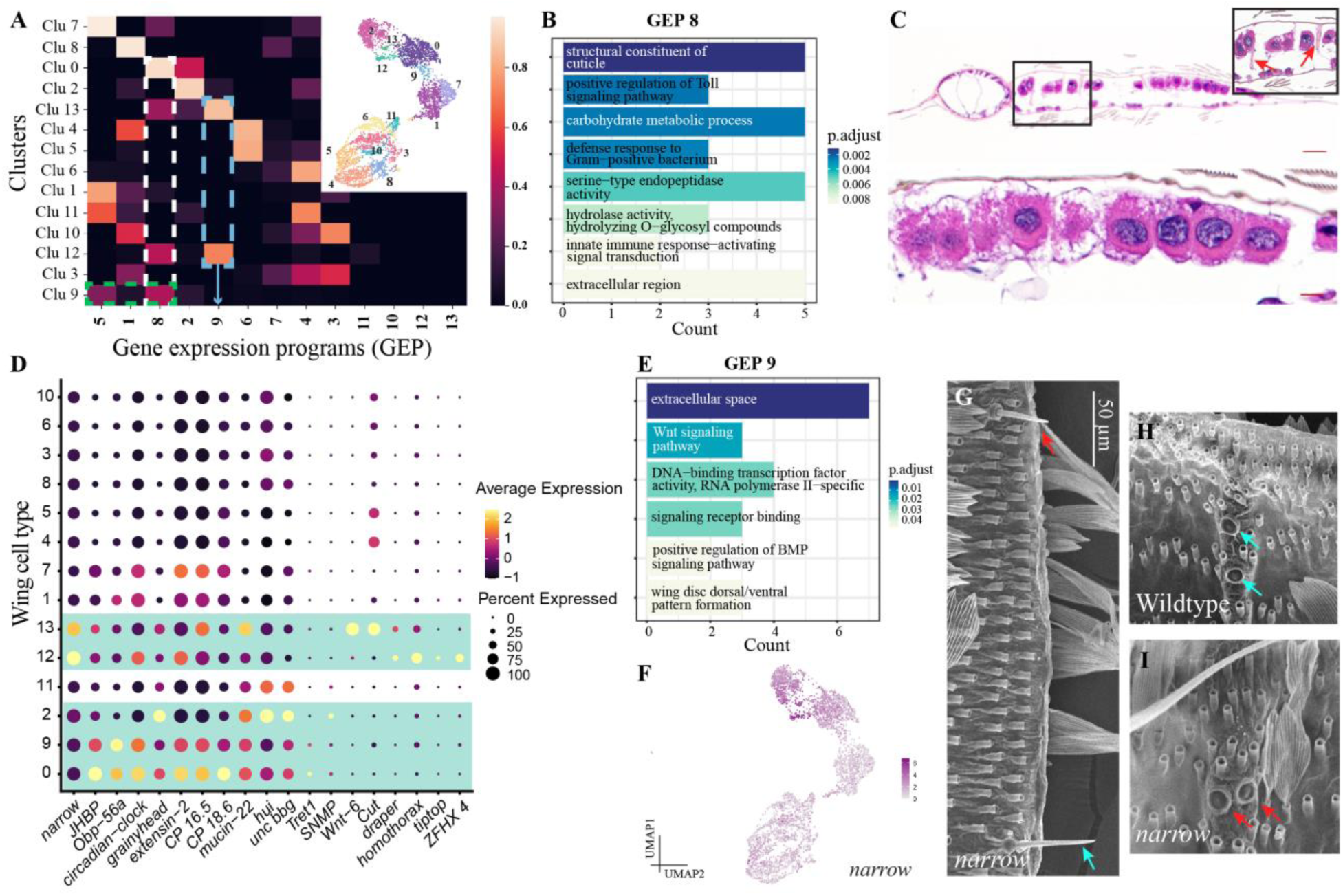
Epidermal and innervated sensory cells on *B. anynana* forewings. (A) GEP usage map highlighting the usage of two different GEPs 8 and 9 across different clusters. Inset shows a cropped UMAP plot of the *B. anynana* single-cell atlas with the corresponding cluster numbers marked. (B) Gene enrichment plot of the top 60 genes within GEP 8. (C) Paraffin embedded cross-sections of the wildtype forewing androconial gland stained with H&E showing large glandular cells restricted to the dorsal surface of the wing, overlaid by cuticle. Pillars of cuticle (red arrows in inset) appear to occur between the two surfaces of the gland that potentially make a reservoir for the pheromones. Scale bars are 10 µm (D) Dotplot of different marker genes highly expressed in clusters 0, 2, 9, 12 and 13. The horizontal green boxes highlight the putative epidermal and innervated macrochaetes on the wing. (E) Gene enrichment plot of the top 60 genes within GEP 9 (F) UMAP expression plot of an uncharacterized gene in *B. anynana* that is homologous to *Drosophila narrow*. (G-I) *narrow* crispant phenotypes affect innervated macrochaete development in *B. anynana* (red arrows). Blue arrows mark wildtype-like marginal mechanosensory bristle in a *narrow* crispant as well as wildtype campaniform sensilla.

To verify the dorsal location of the glandular, pheromone producing cells, we performed paraffin cross-sections of the wildtype gland in males. We indeed found large cells restricted to the dorsal side of the gland, overlaid by cuticle, indicating that cluster 0 comprised of androconia-related, cuticle-synthesizing glandular cells (Fig 5C). To verify this, we knocked out three genes, *grainyhead*, *circadian clock-controlled* gene and *FAR*, all three of which were expressed in cluster 0 (Fig S15A). Though the crispants showed mutations in the targeted genomic regions (Fig S15B), the frequency of indels was low and analysis of pheromone composition from these individuals did not show dramatic variations from the wildtype, potentially because of the low CRISPR efficiency or mistargeting of cells in the small androconial patches (Fig S15C).

Cells in cluster 9 express a combination of GEPs 5 and 8 (Fig 5A, green dashed box) and high levels of both *HR38* and *sens* (Fig 3A, D), and are potentially androconia-related dorsal scale cells.

We annotated cluster 2 as containing tracheal cells or muscle cells. Cells in this cluster strongly expressed GEP 2 which was enriched in genes involved in cell surface, heterophilic cell-cell adhesion, adherans junctions and larval visceral muscle development (Fig S4I). The transcription factor *grainyhead* was also expressed at high levels in clusters 0 and 2 (Fig 5D, S15A). *grainyhead* is known to be important for cuticle formation and epidermal cell development in both vertebrates and invertebrates and in regulating apical cell membrane growth during tracheogenesis (Gangishetti et al., 2012; Hemphälä et al., 2003; Mace et al., 2005; Moussian and Uv, 2005; Yao et al., 2017).

Clusters 12 and 13 were annotated as different types of innervated sensory cells on the wing. These were the smallest clusters, with cluster 12 being *apA-*positive dorsal cells in contrast to the *apA*-negative ventral cluster 13 (Fig 1D). Both clusters shared the unique expression of GEP 9 (Fig 5A dotted blue box) that was enriched for genes involved in Wnt signaling, and dorsal-ventral wing patterning, suggesting that cells in clusters 12 and 13 lie close to the wing margin (Fig 5E). An uncharacterized gene with a C-type lectin domain, homologous to *Drosophila narrow,* was strongly expressed in these two clusters (Fig 5D, F). In addition, cluster 13 expressed *cut* and *wnt-6* at high levels (Fig 5D). Supporting our annotation, strong *cut* expression was seen in regularly spaced cells along the wing margin corresponding to the mechanosensory bristles (Fig S6C, D). High *cut* expression was also seen in the peripheral tissue cells close to the wing margin (Fig S6B) which may have also been captured in cluster 13. Cluster 12 differed from cluster 13 by the expression of a different group of genes such as *homothorax* and *tiptop. homothorax* is expressed along the margin and proximal wing and hinge regions in *Drosophila* and *Heliconius* butterflies (Fig 5D) (Azpiazu and Morata, 2000; Bessa et al., 2009; Hanly et al., 2019). Thus, cells in cluster 12 are probably different sensory cell types close to the wing hinge.

To functionally validate cells in cluster 12 and 13, we first localized the sensory macrochaetes on *B. anynana* pupal wings, using immunostainings with an anti-synapsin antibody, which stains neural synapses, and then knocked out a differentially expressed gene in these clusters. Various kinds of sensory cells were present across the wing disc (Fig S16A-D). Stout mechanosensory bristles occurred on the ventral side of the wing margin (Fig S16A, B - green arrow, E), while dorsally, thin, hair-like bristles were present (Fig S16A, B - blue arrow). These thin sensory organs were also present along the veins and near the base of the wings (Fig S16C – blue arrows) and were different from hair-like, non-innervated scales by their bases, which formed a rotatable structure within the socket (Fig S16D - blue arrow). Campaniform sensilla were also distributed across the wing, along the trachea (Fig S16C, D - red arrows). They occurred in groups at the base of the dorsal wing surface (Fig S16C, G) and in a characteristic pair, one below the other, on the ventral veins near the margin (Fig 5H, S16A - red arrows, S16F). To verify if clusters 12 and 13 indeed correlated with the sensory cell types, we knocked out *narrow* using CRISPR/Cas9 (Fig S16I, L). Like *Drosophila* RNAi mutants, *B. anynana narrow* crispants had distorted wing shapes (Fig S16I) (Ray et al., 2015). Additionally, they exhibited shorter ventral marginal mechanosensory bristles (Fig 5G, S16J, K). In one individual, positioning of the marginal campaniform sensilla was also affected, with the pair now present side by side (Fig 5H, I). This data supports clusters 12 and 13 mapping to various sensory cell types on the different surfaces of the wing.

## Discussion

### Scale cell differentiation is temporally and spatially heterogenous across the wing

In this scRNA-seq analysis of large cells in ∼24 hour *B. anynana* pupal forewings, we annotated putative scale cell clusters and identified scale cells at different stages of their differentiation. At this stage, Cut protein was not expressed in all pIIa cells and in all scale and socket cell pairs. Varying levels of Cut expression were seen in some precursor and divided scales while others did not express Cut at all. It is possible that Cut expression is temporally dynamic and that all scale and socket cell pairs express Cut at later time points. Alternatively, Cut could be marking a particular scale cell type. These hypotheses and the dynamics of Cut expression during butterfly scale development should be tested in future.

### *HR38* and *senseless* are important regulators of scale cell determination and differentiation

We found that *sens* is necessary for both the specification of scale cell precursors and also to specify size, color, and morphology of different scale cell types. Loss of *sens* caused a loss of row-like scale organization, lower scale density, larger scales, the appearance of smaller and scattered silver scales (dorsally) and to most scales becoming yellower. How *sens* determines surface-specific, and scale type-specific variations (cover vs ground, different colors and morphologies) remains unknown. Most likely, it acts in a combinatorial fashion with other transcription factors to determine cell type specificity. In the *Drosophila* peripheral nervous system, *sens* acts as a binary switch during SOP selection, acting downstream of the proneural genes (Jafar-Nejad et al., 2003; Nolo et al., 2000), and later regulating terminal differentiation of the sensory cells (Xie et al., 2007). Loss of *sens* also causes loss of sensory organ precursors via apoptosis (Nolo et al., 2000), potentially explaining the lower scale density phenotype observed in *B. anynana*.

The white eyespot center scales were, however, never affected in *sens* crispants. A possible hypothesis to explain this is the spatial repression of *sens* in the eyespot centers post-transcriptionally via the expression of an eyespot center-specific microRNA (Tian and Monteiro, 2022). A microRNA, *miRNA-9a,* modulates levels of *sens* expression in a dynamic and complex manner in neural precursor cells during *Drosophila* sensory organ development leading to correct SOP specification (Gallicchio et al., 2021; Li et al., 2006). Various such non-coding RNAs were expressed in different cell clusters in our dataset (Supp Fig S7) and future work should investigate their function.

We identified novel functions for *HR38* in scale development. This hormone receptor was necessary for the development of hair-like scales, for the elongation of paddle-like scales, and for regulating their color. *Drosophila* HR38 (DHR38) protein, can form a heterodimeric complex with transactivated Ultraspiracles (USP) protein to mediate an atypical ecdysteroid signaling pathway, independent of that transduced by the Ecdysone receptor/USP heterodimer (Baker et al., 2003). Mutations in *DHR38* lead to flies with abnormal cuticle formation (Kozlova et al., 1998) and reduced levels of expression of various cuticle genes (Kozlova et al., 2009). *HR38* has also been implicated in regulating the expression of genes in the melanin pathway (Kalay et al., 2016; Sekine et al., 2011). Together, these results suggest that ecdysteroid hormonal regulation via *HR38* plays a role in lepidopteran scale cell type development.

### Pheromone producing glandular cells are modified dorsal epithelial cells

We annotated dorsal-specific cells in our dataset with high metabolism as pheromone-producing glandular cells. These cells expressed high levels of cuticle producing genes as well as an odorant binding protein, Obp-56a. Odorant binding proteins are highly expressed in pheromone glands (Zhang et al., 2015), and Obp-56a binds to fatty acids such as palmitic acid and has been proposed as a solubilizer of fatty acids during digestion in blowflies (Ishida et al., 2013). Palmitic acid is a precursor of *B. anynana* sex pheromones (Liénard et al., 2014), and thus Obp- 56a may be involved in solubilization and transport of sex pheromones in the gland cells.

Further, though our analysis of pheromone composition in *grainyhead*, *circadian clock-controlled* and *FAR* crispants didn’t identify any changes in levels of both MSPs 1 and 3, the mosaic nature of CRISPR experiments might have failed to target the small number of glandular cells. Alternatively, genes such as *grainyhead* and *circadian clock-controlled* (which has a juvenile hormone binding protein domain) may function in cuticle production or transport and temporal determination of pheromone production which was not captured in our analysis. In future, creating mutant lines would be a more precise way of investigating the roles of these genes in gland and pheromone development.

### Scale cell types and a model for scale type differentiation

Butterfly wings are covered by a diverse array of scale cells that have largely been classified into a two-layer system consisting of large cover scales overlying smaller ground scales. In our study, we came across an intermediate scale cell type, distinct from the cover and ground scales. Though mentions of such intermediate scale types have been made in previous literature from moths (Kristensen, 2012), its occurrence in butterflies had not been noted before. The presence of this intermediate scale type, which varies across and within wing surfaces, hints at the yet uncharacterized complexity of scale cell types on butterfly wings, and their developmental programs. Therefore, the first major challenge to be able to understand the evolution and development of scale cell diversity in the future, would be to systematically classify different scale types. Borrowing from concepts in neuronal cell type classification and taxonomic principles, this would entail using multiple criteria (e.g., morphology, color, and molecular basis in the case of scales), adopting a hierarchical classification system to account for relationships between the different types and initially classifying types within specific wing regions such as dorsal vs. ventral (Zeng and Sanes, 2017).

The generation of diverse scale cell types on butterfly wings will likely involve processes known to generate other diverse cell types such as neurons or sensory organs (Arendt, 2008; Klann et al., 2021). In most cases a core differentiation program, consisting of actin elongation and chitin synthesis, is modified by spatial or temporal control of regulatory factors, outside the core, that specify different scale cell fates. Genes such as *Pax5* (and perhaps also *ASH2)* might represent top-level regulators of the core scale cell program. Once specified, genes such as *sens* and *HR38*, appear to direct cell type specific differentiation. For example, low levels of *HR38* might be required to differentiate hair-like scales, whereas *sens* might be more highly expressed in cover scales, to make these larger than ground scales. These hypotheses require future testing.

Our analysis, done on large cells on the wings at ∼24 hours after pupation, identified many clusters of dorsal and ventral scale cells. However, though we annotated these clusters broadly as scale cells and functionally verified this annotation with a few highly expressed marker genes, we were unable to further annotate individual clusters, with markers for scale color, for instance. At this early pupal stage, scale cells diverged more strongly in their transcriptomic signals for genes involved in developmental progression coupled with broad morphological and functional diversity of machrochaeta, rather than genes involved in color production per se. Furthermore, previously investigated transcription factors that are already marking the different colored scales at early stages of development (e.g., *spalt*, *Distal-less*, *engrailed*, *optix*, *doublesex,* etc) (Banerjee et al., 2020; Brunetti et al., 2001; Prakash and Monteiro, 2020b), were not readily picked up by the single-cell sequencing, perhaps due to low levels of expression. To further annotate and distinguish various scale cell types, later stages of development could be sampled alongside increased depth of sequencing. Overall, our results provide a foundational atlas for future explorations of the genetic, molecular, evolutionary, and developmental basis of the various cell types in butterfly wings.

## Materials and methods

### Butterfly husbandry

*Bicyclus anynana* butterflies were reared in a temperature-controlled room at 27°C, 65% humidity and a 12:12 hour light:dark cycle. Caterpillars were fed on corn plants and adults were fed on mashed bananas.

### Preparation of a single cell suspension, library preparation and sequencing

The preparation of a single-cell suspension from 22.5-25-hour male pupal forewings of *B. anynana* butterflies has been documented in-depth in a protocol paper (Prakash and Monteiro, 2020a). Briefly, 14 male pupal forewings were dissected into multiple tubes with sterile PBS, washed, then dissociated in warm 5X TrypLE with continuous trituration for a total of 15-20 minutes. The dissociated cell mixture was filtered with a 41 µm filter and collected by centrifuging at 300 g, 4 °C for 5 minutes. Samples were pooled and gently resuspended into 1mL of cold 1X PBS + 0.01% BSA. Cells were counted in a Countess Automated cell counter by mixing 5 µl of sample with 5 µl of 0.4% trypan blue and loading it into a Countess cell counting chamber. Before going for fluorescence-activated cell sorting (FACS), 5 µl of a 300 µM intermediate concentration of DAPI (Invitrogen by Thermo Fisher Scientific, Cat. no.: D1306) was added per mL of sample. Cells larger than 6 µm in diameter were sorted using FACS and collected in a 1.5 mL LoBind Eppendorf tube pre-filled with 50 µl of cold 1X PBS + 0.01% BSA. Around 104,000 cells were collected and precipitated by centrifuging at 900g, 4 °C for 3 minutes. Cells were resuspended in 30 µl of cold 1X PBS + 0.01% BSA, with a final cell count of 1650 cells/µl and 88% viability.

Cells were processed via the 10X Genomics microfluidics system at the Genome Institute of Singapore. Single-cell libraries were generated using the Chromium Single cell 3’ Library Kit V3 with a targeted cell recovery of 7000 cells. The quality of the generated cDNA library was quantified using an Agilent Bioanlyzer and sequenced on one lane of HiSeq4K 2x151bp (multiplex) at the Genome Institute of Singapore.

### Data pre-processing and quality control

Barcode and UMI trimming, demultiplexing, and quality assessment of the reads was performed using the scPipe R package (Tian et al., 2018). The workflow was as follows: The FASTQ files containing the reads were first reformatted to trim the barcodes and UMI sequences and move it to the header. Reads were then aligned to the *Bicyclus anynana* genome (v1.2) and mapped to exons using an annotation file - Bicyclus_anynana_v1.2.gff3 downloaded from Lepbase (Challi et al., 2016). Demultiplexing was done based on barcodes. Of the 346,685,500 initial reads, 72.36% had a barcode match. The generated gene count matrix consisted of 13631 genes and 9986 cells, with a mean count per cell of ∼10700 reads and mean number of genes per cell of ∼1400.

We next discarded low-quality cells, empty droplets with no cells, and potential doublets from the gene count matrix. Cells with counts per cell>100000 were removed and genes with less than 10 counts summed across all cells were discarded. The DropletUtils R package (Griffiths et al., 2018; Lun et al., 2019) was used to filter our empty droplets using a FDR <=0.001. The detect_outlier function in scPipe was then used to detect outliers (both low quality cells and potential doublets) using a Gaussian mixture model. Additionally, we removed 18 histone and 53 ribosomal protein genes because an initial analysis of the data indicated unsupervised clustering based on these genes, masking underlying patterns (Table S2). This led to a gene count matrix of 5268 cells and 10743 genes used in the initial exploratory analysis detailed below. A Seurat object (Stuart et al., 2019) was created using the count matrix with min.cells = 5 i.e., keeping features that were present in at least 5 cells. The count matrix for the Seurat analysis comprised 5268 cells and 10696 genes.

### Analysis of scRNAseq data, unsupervised clustering and annotation of clusters

Downstream analysis was carried out in two ways in R v4.3.1 (R Core Team, 2021): using Seurat v3.1 (Stuart et al., 2019) (https://satijalab.org/seurat/articles/pbmc3k_tutorial.html) or scran (Lun et al., 2016) (https://bioconductor.org/packages/devel/bioc/vignettes/scran/inst/doc/scran.html). Initial data exploration was carried out using scran. First, scaling normalization was performed using a deconvolution strategy with the quickCluster, computeSumFactors and logNormCounts functions. Highly variable genes were identified by variance modelling using the modelGenVarByPoisson function and getTopHVGs function was used to select the top 50% (prop = 0.5; var.field = “bio”) of genes. The reduced data matrix with the highly variable genes was used as an input for a Principal Component Analysis (PCA). The number of principal components to retain in the analysis was computed using getClusteredPCs function. A shared nearest neighbor graph was constructed using buildSNNGraph function, following which, clusters were identified using the igraph package. Varying values of k changed cluster resolution with smaller k values giving finer clusters. Different k values were used for the initial exploration of the data. runUMAP and plotUMAP functions helped visualize the clusters.

In the Seurat analysis, transcript counts for each cell were log normalized using the NormalizeData function. The top 1000 highly variable transcripts across all cells were selected using the FindVariableFeatures function and ‘vst’ as the selection method. Transcript expressions across cells were then centered and scaled using the ScaleData function. Dimensionality reduction was done using PCA on the scaled data matrix with the previously determined variable features. Thereafter, FindNeighbors function was used to construct a shared nearest neighbor graph using the first 25 principal components and prune.SNN = 1/15. Clustering was performed using the FindClusters function and a resolution of 0.8. The wing cell type atlas was generated using the RunUMAP function on the first 50 principal components and plotted using DimPlot.

Clusters were annotated using a combination of different methods. Initially, expression of known marker genes from the *Drosophia melanogaster* macrochaete development literature were visualized using the FeaturePlot function. In parallel, genes highly and differentially expressed between various clusters were identified from both the Seurat and scran analyses using FindAllMarkers (Seurat), FindMarkers (Seurat) or findMarkers (scran) functions.

### Consensus non-Negative Matrix Factoriztion (cNMF)

cNMF was run on the cells-by-transcripts count matrix (5268 cells by 10743 transcripts) to identify gene expression programs (Kotliar et al., 2019). The steps followed were based on the code provided in (Brückner et al., 2021). We used the 2000 most over-dispersed transcripts for the factorizations and ran 200 replicates of NMFs for each value of K ranging from 5 to 20. Replicates for each K were then combined to obtain a consensus, and this was used to estimate the stability and error for each K. The stability-error plot as a function of K was used to select the best choice of K for our analysis. Though K=10 had the maximum stability solution, we found that this K value provided poorer resolution of our clusters in the downstream analysis. We chose to go for K=13 which had the second highest stability vs error. We re-ran the consensus with K=13 and a density threshold of 0.18, to obtain the consensus GEP usage scores for each cell. The normalized GEP usage scores were combined with the cluster annotations for each cell obtained from Seurat to calculate the proportional GEP usage in each cluster, visualized as a heatmap.

Gene scores were extracted from the generated gene_spectra_score text file and the top 60 enriched genes in each GEP were obtained. Gene enrichment analysis was carried out using the enricher function (pvalueCutoff and qvalueCutoff = 0.05, pAdjustMethod = “BH”) from the clusterProfiler package (Yu et al., 2012).

### Immunostainings

We used the 2B10 mouse anti-Cut primary antibody raised against a *Drosophila* Cut antigen. 2B10 was deposited to the DSHB by Rubin, G.M. (DSHB Hybridoma Product 2B10). This primary antibody has been previously used to target Cut protein expression in many Lepidopteran species (Macdonald et al., 2010).The mouse 3C11 (anti-SYNORF1) was used to target synapsin. 3C11 (anti SYNORF1) was deposited to the DSHB by Buchner, E. (DSHB Hybridoma Product 3C11 (anti SYNORF1)). An Alexa Fluor 488-conjugated goat anti-mouse antibody (Jackson ImmunoResearch Laboratories, Inc.) was used as a secondary antibody.

Pupal wing tissues at various time points were dissected and fixed in fix buffer (0.1M PIPES pH 6.9, 1mM EGTA pH 6.9, 1% Triton X-100, 2mM MgSO4) at room temperature. Fixation was done for 30 min following the addition of formaldehyde (final concentration of 4% directly to the wells). The wings were then moved onto ice, washed five times with PBS and transferred to block buffer (50mM 626 Tris pH 6.8, 150mM NaCl, 0.5% IGEPAL, 1mg/ml BSA) at 4 °C overnight. Incubation with primary antibody at a concentration of 2-2.5 µg/ml in wash buffer (50mM Tris pH 6.8, 150mM NaCl, 0.5% IGEPAL, 1mg/ml BSA) was carried out at room temperature for 1 hour, followed by four washes with wash buffer. Wings were then incubated with secondary antibody (1:500) for 30 minutes at room temperature followed by 5-10 washes to remove the secondary antibody. Wing tissues were counterstained with DAPI (1:100) for 5-10 minutes, followed by further washing. Control stains used only secondary antibodies. Mounted wings were imaged on an Olympus FLUOVIEW FV3000 confocal microscope.

For staining wings with CellMask Orange Plasma membrane stain (Invitrogen, Cat no. C10045) and DAPI, wings were dissected and immediately placed in PBS with the CellMask Orange stain (1:300) for 10 min at room temperature. This was because the CellMask Plasma membrane stain does not survive permeabilization. The stain was then removed, and wings were fixed in 4% formaldehyde in PBS at room temperature for 20 min, followed by 5 washes with cold PBS. Tissues were counterstained with DAPI (1:100) for 5-10 minutes, then washed with PBS 3-5 times. Mounted wings were imaged on an Olympus FLUOVIEW FV3000 confocal microscope.

### Optical imaging and Scanning Electron Microscopy

Optical images of single scales and wildtype or crispant wings were captured using a Leica DMS 1000 microscope.

For scanning electron microscopy, desired regions of the wildtype or crispant adult wings were cut using scissors and mounted onto carbon tape. For imaging different sensory cell types, the scales were carefully and gently brushed off the wings using a paintbrush. Wing pieces were sputter coated with gold using a JFC-1100 Fine Coat Ion Sputter (JEOL Ltd. Japan) and imaged on a JEOL JSM-6510LV scanning electron microscope (JEOL Ltd. Japan) located at the Center for Bioimaging Sciences (CBIS, NUS).

### Phylogenetic analysis

To identify the evolutionary relatedness of the *Bicyclus anynana* AS-C homolog, identified in the scRNA-seq analysis, to other Lepidopteran, Dipteran and Coleopteran AS-C genes, we compiled a list of protein sequences of AS-C homologs from NCBI, based on (Zhou et al., 2008). The final dataset consisted of 12 amino acid sequences with a total of 503 positions. Evolutionary analyses were conducted in MEGA X (Kumar et al., 2018). The evolutionary history was inferred by using the Maximum Likelihood method and JTT matrix-based model (Jones et al., 1992). Initial tree(s) for the heuristic search were obtained automatically by applying Neighbor-Join and BioNJ algorithms to a matrix of pairwise distances estimated using a JTT model, and then selecting the topology with superior log likelihood value. The tree with the highest log likelihood (-7192.28) was selected.

### Probe synthesis and *in situ* hybridization

mRNA sequences were amplified from cDNA using the primers specified in Supplementary Table S3 and cloned into a PGEM-T Easy vector (Promega). Colonies containing plasmids with inserts of the right size were selected using blue-white screening. Purified plasmids were used to prepare digoxigenin-labelled sense and anti-sense riboprobes using T7 and SP6 RNA polymerases (Roche). Probes were purified using ethanol precipitation and resuspended in 1:1 volume of DEPC-treated water:formamide.

Pupal wings at various time points were dissected and placed in glass well plates containing PBST (PBS+0.1% Tween20). Tissues were fixed at room temperature for 30 min in 4% formaldehyde in PBST followed by 3 washes in cold PBST. The wings were then incubated in 25 μg/mL proteinase K in cold PBST for 5 minutes, washed twice with 2 mg/mL glycine in cold PBST, then 5 times with cold PBST before being gradually transferred to a prehybridization buffer (5X saline sodium citrate (pH 4.5), 50% formamide, 0.1% Tween20 and 100 μg/mL denatured salmon sperm DNA). Tissues in prehybridization buffer were incubated at 60°C for 1 hour then transferred to a hybridization buffer (prehybridization buffer with 1 mg/mL glycine and 180 ng/ml DIG labelled riboprobe) overnight. This was followed by 5-6 rounds of washes with prehybridization buffer at 60°C, a gradual transfer back to PBST at room temperature, 3 washes in PBST and blocking overnight at 4°C (PBST + 1% BSA). Probe detection was by tissue incubation in 1:3000 anti-DIG alkaline phosphatase (Roche) in block buffer for 1 hour, 2 washes in alkaline phosphatase buffer (100 mM Tris (pH 9.5), 100 mM NaCl, 5 mM MgCl2, 0.1% Tween) and a final incubation at room temperature in NBT/BCIP solution (Promega) till color developed. The reaction was stopped by washing in PBST. Wings were mounted on slides and imaged on a Leica DMS 1000 microscope. High magnification images were captured using a 100X lens of a uSight-2000-Ni microspectrophotometer (Technospex Pte. Ltd., Singapore) and a Touptek U3CMOS-05 camera.

### CRISPR-Cas9 gene editing

CRISPR-Cas9 gene editing was done following previously published protocols (Banerjee and Monteiro, 2018). Guide RNAs targeting genes of interest, preferably protein domains, were manually designed by searching for GGN18NGG sequences (Table S4). Injection mixtures containing 500 ng/µl of Cas 9 protein (NEB; Cat. no.: M0641) and 300 ng/µl of guide RNA were prepared and injected into eggs within 4 hours of egg laying. Caterpillars were reared on corn plants and adults were scored for their phenotypes.

### Genotyping crispants

Genomic DNA from thoracic tissue and legs of potential crispant individuals was isolated using the Kaneka Easy DNA Extraction Kit 2 (Cat. No. KN-T110005, Kaneka, Japan). Next generation sequencing was used to verify the CRISPR edits. Indexed libraries were prepared in a two-step PCR process. First, targeted regions were amplified using gene specific primers appended to reading primers: Forward: 5’ACACTCTTTCCCTACACGACGCTCTTCCGATCT3’; Reverse: 5’GTGACTGGAGTTCAGACGTGTGCTCTTCCGATCT3’ (Table S5). Gene of interest amplicon size was kept to <300 bp. In the second PCR step, indices and Illumina adapter sequences were appended. The Illumina adapters D501-D508 and D701-D712 were used to multiplex samples of different genes and different individuals together. PCR products were then purified using ethanol purification, pooled together and sequenced on an iSeq 100 (2 x 150 bp) at the Chae lab, National University of Singapore. Demultiplexed FASTQ files were visualized online using the web-tool “CRISPR-RGEN Tools” (http://www.rgenome.net/).

### Histology

Androconial glands from the forewings of adult male butterflies were cut from the wings using a dissection scissors and descaled using a brush. Tissue pieces were fixed in 4% formaldehyde in PBS overnight. Samples were then passed to the Advanced Molecular Pathology Laboratory (AMPL) at the A*STAR Institute of Molecular and Cell Biology, Biopolis, Singapore for downstream processing. Tissues were embedded in paraffin blocks, sectioned and stained with H&E. Sections were imaged using the imaging system on a uSight-2000-Ni microspectrophotometer (Technospex Pte. Ltd., Singapore) and a Touptek U3CMOS-05 camera.

### Sex pheromone composition and quantification

The composition of the sex pheromone blend and the amount of the two components (called MSP1 and MSP3 for Male Sex Pheromone) were compared between wildtype and crispant individuals using gas chromatography – mass spectrometry. Three days-old crispant butterflies were anesthetized at -80°C for 5 minutes. The forewing was cut at the base of the thorax with fine scissors and placed for 15 minutes in a glass vial containing 500uL of hexane. 400uL of the solution containing the dissolved sex pheromones was extracted to a new vial, to which 10 µg/mL of methyl stearate (Merck, Singapore) was added as an internal standard. All tools and vials were pre-rinsed with hexane, and all extractions were done at 2 pm to prevent the effects of daily fluctuations in pheromone titers.

The extracts were analyzed with a Shimadzu Gas-Chromatography-QQQ Mass Spectrometer equipped with a DB-5 column, using the following set up: electron ionization (EI) was done at 70 eV, 1 μL of each extract was injected splitless, with the injector temperature set up at 250 °C. Helium was used as carrier gas, with the flow set at 1.9 mL/min. The column temperature gradient began at 50 °C, increased to 210 °C at a rate of 35 °C/min, then increased to 280 °C at a rate of 3 °C/min. The detector was set to unit mass resolution and 3 scans/sec, from m/z 37 to 500. Chromatograms and mass spectra were analyzed using the Shimadzu GCMSsolution software v. 4.11. The components and precursors are well documented and were identified based on their known mass spectrum and retention time (Dion et al., 2016; Liénard et al., 2014; Nieberding et al., 2008). The amount of sex pheromone components was calculated by normalizing the area of the component peak to the peak of the internal standard.

## Acknowledgements

We thank Dr. Rajaganesan Ramaswamy for helping with the code to run cNMF. We acknowledge Tong Yan and Chong Ping Lee from the NUS Centre for Bioimaging Sciences for their technical support in using the confocal and scanning electron microscopes. We thank the Advanced Molecular Pathology Laboratory at A*Star’s Institute of Molecular and Cell Biology, Singapore for their service in embedding and sectioning wing paraffin samples. We are grateful to Yi Yun and the Eunyoung Chae Lab at the Department of Biological Sciences, NUS for the amplicon library preparation protocols and the iSeq runs. We thank the NUS Environmental Research Institute for the use of the GC-QQQ and associated equipment. We are grateful to Greenology for providing corn plants.

## Author contributions

Conceptualization, A.P. and A.M.; Methodology, A.P., E.D.; Formal Analysis, A.P.; Investigation, A.P., E.D.; Writing – Original Draft, A.P.; Writing – Review & Editing, A.P., E.D., A.M.; Visualization, A.P.; Funding Acquisition, A.P. and A.M.; Supervision, A.M.

## Data Availability

Raw sequencing reads and processed matrix files will be available through NCBI GEO (Accession Number: TBA).

## References

Aiello, B.R., Stanchak, K.E., Weber, A.I., Deora, T., Sponberg, S., and Brunton, B.W. (2021). Spatial distribution of campaniform sensilla mechanosensors on wings: form, function, and phylogeny. Curr. Opin. Insect Sci. 48, 8–17. 10.1016/j.cois.2021.06.002.

Arendt, D. (2008). The evolution of cell types in animals: Emerging principles from molecular studies. Nat. Rev. Genet. 9, 868–882. 10.1038/nrg2416.

Artavanis-Tsakonas, S., and Simpson, P. (1991). Choosing a cell fate: a view from the Notch locus. Trends Genet. 7, 403–408. 10.1016/0168-9525(91)90264-q.

Ashour, D.J., Durney, C.H., Planelles-Herrero, V.J., Stevens, T.J., Feng, J.J., and Röper, K. (2022). Zasp52 strengthens whole embryo tissue integrity through supracellular actomyosin networks. BioRxiv 10.1101/2022.10.11.511783.

Azpiazu, N., and Morata, G. (2000). Function and regulation of homothorax in the wing imaginal disc of Drosophila. Development 127, 2685–2693. 10.1242/dev.127.12.2685.

Baker, K.D., Shewchuk, L.M., Kozlova, T., Makishima, M., Hassell, A., Wisely, B., Caravella, J.A., Lambert, M.H., Reinking, J.L., Krause, H., et al. (2003). The Drosophila orphan nuclear receptor DHR38 mediates an atypical ecdysteroid signaling pathway. Cell 113, 731–742. 10.1016/S0092-8674(03)00420-3.

Banerjee, T. Das, and Monteiro, A. (2018). Crispr-cas9 mediated genome editing in bicyclus anynana butterflies.

Banerjee, T. Das, Ramos, D., and Monteiro, A. (2020). Expression of Multiple engrailed Family Genes in Eyespots of Bicyclus anynana Butterflies Does Not Implicate the Duplication Events in the Evolution of This Morphological Novelty. Front. Ecol. Evol. 8. 10.3389/fevo.2020.00227.

Bessa, J., Carmona, L., and Casares, F. (2009). Zinc-finger paralogues tsh and tio are functionally equivalent during imaginal development in Drosophila and maintain their expression levels through auto- and cross-negative feedback loops. Dev. Dyn. 238, 19–28. 10.1002/dvdy.21808.

Brückner, A., Badroos, J.M., Learsch, R.W., Yousefelahiyeh, M., Kitchen, S.A., and Parker, J. (2021). Evolutionary assembly of cooperating cell types in an animal chemical defense system. Cell 184, 6138–6156.e28. 10.1016/j.cell.2021.11.014.

Brunetti, C.R., Selegue, J.E., Monteiro, A., French, V., Brakefield, P.M., and Carroll, S.B. (2001). The generation and diversification of butterfly eyespot color patterns. Curr. Biol. 11, 1578–1585. .

Challi, R.J., Kumar, S., Dasmahapatra, K.K., Jiggins, C.D., and Blaxter, M. (2016). Lepbase: the Lepidopteran genome database. BioRxiv 10.1101/056994.

Dickerson, B.H., Aldworth, Z.N., and Daniel, T.L. (2014). Control of moth flight posture is mediated by wing mechanosensory feedback. J. Exp. Biol. 217, 2301–2308. 10.1242/jeb.103770.

Dinwiddie, A., Null, R., Pizzano, M., Chuong, L., Leigh Krup, A., Ee Tan, H., and Patel, N.H. (2014). Dynamics of F-actin prefigure the structure of butterfly wing scales. Dev. Biol. 392, 404–418. 10.1016/j.ydbio.2014.06.005.

Dion, E., Monteiro, A., and Yew, J.Y. (2016). Phenotypic plasticity in sex pheromone production in Bicyclus anynana butterflies. Sci. Rep. 6, 1–13. 10.1038/srep39002.

Fabian, J., Siwanowicz, I., Uhrhan, M., Maeda, M., Bomphrey, R.J., and Lin, H.T. (2022). Systematic characterization of wing mechanosensors that monitor airflow and wing deformations. IScience 25, 104150. 10.1016/j.isci.2022.104150.

Ficarrotta, V., Hanly, J.J., Loh, L.S., Francescutti, C.M., Ren, A., Tunström, K., Wheat, C.W., Porter, A.H., Counterman, B.A., and Martin, A. (2022). A genetic switch for male UV iridescence in an incipient species pair of sulphur butterflies. Proc. Natl. Acad. Sci. U. S. A. 119. 10.1073/pnas.2109255118.

Furman, D., and Bukharina, T. (2008). How Drosophila melanogaster Forms its Mechanoreceptors. Curr. Genomics 9, 312–323. 10.2174/138920208785133271.

Galant, R., Skeath, J.B., Paddock, S., Lewis, D.L., and Carroll, S.B. (1998). Expression pattern of a butterfly achaete-scute homolog reveals the homology of butterfly wing scales and insect sensory bristles. Curr. Biol. 8, 807–813. 10.1016/S0960-9822(98)70322-7.

Gallicchio, L., Griffiths-Jones, S., and Ronshaugen, M. (2021). Single-cell visualization of miR-9a and Senseless co-expression during Drosophila melanogaster embryonic and larval peripheral nervous system development. G3 Genes, Genomes, Genet. 11, 1–11. 10.1093/G3JOURNAL/JKAA010.

Gangishetti, U., Veerkamp, J., Bezdan, D., Schwarz, H., Lohmann, I., and Moussian, B. (2012). The transcription factor Grainy head and the steroid hormone ecdysone cooperate during differentiation of the skin of Drosophila melanogaster. Insect Mol. Biol. 21, 283–295. 10.1111/j.1365-2583.2012.01134.x.

Ghiradella, H. (1991). Light and color on the wing : butterflies and moths structural colors in. Appl. Opt. 30, 3492–3500. 10.1364/AO.30.003492.

Greenstein, M.E. (1972). The ultrastructure of developing wings in the giant silkmoth, Hyalophora cecropia. II. Scale-forming and socket-forming cells. J. Morphol. 136, 23–51. 10.1002/jmor.1051360103.

Griffiths, J.A., Richard, A.C., Bach, K., Lun, A.T.L., and Marioni, J.C. (2018). Detection and removal of barcode swapping in single-cell RNA-seq data. Nat. Commun. 9, 2667. 10.1038/s41467-018-05083-x.

Hanly, J.J., Wallbank, R.W.R., McMillan, W.O., and Jiggins, C.D. (2019). Conservation and flexibility in the gene regulatory landscape of heliconiine butterfly wings. Evodevo 10, 1–14. 10.1186/s13227-019-0127-4.

Hartenstein, V., and Posakony, J.W. (1989). Development of adult sensilla on the wing and notum of Drosophila melanogaster. Development 107, 389–405. 10.1242/dev.107.2.389.

Hartenstein, V., and Posakony, J.W. (1990). A dual function of the Notch gene in Drosophila sensillum development. Dev. Biol. 142, 13–30. 10.1016/0012-1606(90)90147-b.

Hemphälä, J., Uv, A., Cantera, R., Bray, S., and Samakovlis, C. (2003). Grainy head controls apical membrane growth and tube elongation in response to Branchless/FGF signalling. Development 130, 249–258. 10.1242/dev.00218.

Ishida, Y., Ishibashi, J., and Leal, W.S. (2013). Fatty Acid Solubilizer from the Oral Disk of the Blowfly. PLoS One 8, 1–9. 10.1371/journal.pone.0051779.

Jafar-Nejad, H., Acar, M., Nolo, R., Lacin, H., Pan, H., Parkhurst, S.M., and Bellen, H.J. (2003). Senseless acts as a binary switch during sensory organ precursor selection. Genes Dev. 17, 2966–2978. 10.1101/gad.1122403.

Jones, D.T., Taylor, W.R., and Thornton, J.M. (1992). The rapid generation of mutation data matrices from protein sequences. Comput. Appl. Biosci. 8, 275–282. 10.1093/bioinformatics/8.3.275.

Kalay, G., Lusk, R., Dome, M., Hens, K., Deplancke, B., and Wittkopp, P.J. (2016). Potential Direct Regulators of the Drosophila yellow Gene Identified by Yeast One-Hybrid and RNAi Screens. G3 Genes|Genomes|Genetics 6, 3419–3430. 10.1534/g3.116.032607.

Kavaler, J., Fu, W., Duan, H., Noll, M., and Posakony, J.W. (1999). An essential role for the Drosophila Pax2 homolog in the differentiation of adult sensory organs. Development 126, 2261–2272. 10.1242/dev.126.10.2261.

Klann, M., Schacht, M.I., Benton, M.A., and Stollewerk, A. (2021). Functional analysis of sense organ specification in the Tribolium castaneum larva reveals divergent mechanisms in insects. BMC Biol. 19, 1–21. 10.1186/s12915-021-00948-y.

Kotliar, D., Veres, A., Nagy, M.A., Tabrizi, S., Hodis, E., Melton, D.A., and Sabeti, P.C. (2019). Identifying gene expression programs of cell-type identity and cellular activity with single-cell RNA-Seq. Elife 8, 1–26. 10.7554/eLife.43803.

Kozlova, T., Pokholkova, G. V., Tzertzinis, G., Sutherland, J.D., Zhimulev, I.F., and Kafatos, F.C. (1998). Drosophila hormone receptor 38 functions in metamorphosis: A role in adult cuticle formation. Genetics 149, 1465–1475. 10.1093/genetics/149.3.1465.

Kozlova, T., Lam, G., and Thummel, C.S. (2009). Drosophila DHR38 nuclear receptor is required for adult cuticle integrity at eclosion. Dev. Dyn. 238, 701–707. 10.1002/dvdy.21860.

Kristensen, N. (2012). Lepidoptera, Moths and Butterflies: Vol 2 Morphology, Physiology, and Development (De Gruyter).

Kumar, S., Stecher, G., Li, M., Knyaz, C., and Tamura, K. (2018). MEGA X: Molecular Evolutionary Genetics Analysis across Computing Platforms. Mol. Biol. Evol. 35, 1547–1549. 10.1093/molbev/msy096.

Lees, A.D., Waddington, C.H., and Gray, J. (1942). The development of the bristles in normal and some mutant types of *Drosophila melanogaster*. Proc. R. Soc. London. Ser. B - Biol. Sci. 131, 87–110. 10.1098/rspb.1942.0019.

Li, Y., Wang, F., Lee, J.A., and Gao, F.B. (2006). MicroRNA-9a ensures the precise specification of sensory organ precursors in Drosophila. Genes Dev. 20, 2793–2805. 10.1101/gad.1466306.

Liénard, M.A., Wang, H.L., Lassance, J.M., and Löfstedt, C. (2014). Sex pheromone biosynthetic pathways are conserved between moths and the butterfly Bicyclus anynana. Nat. Commun. 5. 10.1038/ncomms4957.

Lun, A.T.L., McCarthy, D.J., and Marioni, J.C. (2016). A step-by-step workflow for low-level analysis of single-cell RNA-seq data with Bioconductor. F1000Research 5, 2122. 10.12688/f1000research.9501.2.

Lun, A.T.L., Riesenfeld, S., Andrews, T., Dao, T.P., Gomes, T., Marioni, J.C., and Jamboree, participants in the 1st H.C.A. (2019). EmptyDrops: distinguishing cells from empty droplets in droplet-based single-cell RNA sequencing data. Genome Biol. 20, 63. 10.1186/s13059-019-1662-y.

Macdonald, W.P., Martin, A., and Reed, R.D. (2010). Butterfly wings shaped by a molecular cookie cutter: Evolutionary radiation of lepidopteran wing shapes associated with a derived Cut/wingless wing margin boundary system. Evol. Dev. 12, 296–304. 10.1111/j.1525-142X.2010.00415.x.

Mace, K.A., Pearson, J.C., and McGinnis, W. (2005). An Epidermal Barrier Wound Repair Pathway in Drosophila Is Mediated by grainy head. Science (80-.). 308, 381–385. 10.1126/science.1107573.

Miller, S.W., Avidor-Reiss, T., Polyanovsky, A., and Posakony, J.W. (2009). Complex interplay of three transcription factors in controlling the tormogen differentiation program of Drosophila mechanoreceptors. Dev. Biol. 329, 386–399. 10.1016/j.ydbio.2009.02.009.

Moussian, B., and Uv, A.E. (2005). An ancient control of epithelial barrier formation and wound healing. BioEssays 27, 987–990. 10.1002/bies.20308.

Nieberding, C.M., de Vos, H., Schneider, M. V., Lassance, J.M., Estramil, N., Andersson, J., Bång, J., Hedenström, E., Löfstedt, C., and Brakefield, P.M. (2008). The male sex pheromone of the butterfly Bicyclus anynana: Towards an evolutionary analysis. PLoS One 3, 1–12. 10.1371/journal.pone.0002751.

Nolo, R., Abbott, L.A., and Bellen, H.J. (2000). Senseless, a Zn finger transcription factor, is necessary and sufficient for sensory organ development in Drosophila. Cell 102, 349–362. 10.1016/S0092-8674(00)00040-4.

Overton, J. (1966). Microtubules and microfibrils in morphogenesis of the scale cells of Ephestia kühniella. J. Cell Biol. 29, 293–305. 10.1083/jcb.29.2.293.

Prakash, A., and Monteiro, A. (2018). *apterous A* specifies dorsal wing patterns and sexual traits in butterflies. Proc. R. Soc. B Biol. Sci. 10.1098/rspb.2017.2685.

Prakash, A., and Monteiro, A. (2020a). Cell dissociation from butterfly pupal wing tissues for single-cell RNA sequencing. Methods Protoc. 3. 10.3390/mps3040072.

Prakash, A., and Monteiro, A. (2020b). Doublesex Mediates the Development of Sex-Specific Pheromone Organs in Bicyclus Butterflies via Multiple Mechanisms. Mol. Biol. Evol. 37, 1694– 1707. 10.1093/molbev/msaa039.

Quinlan, M.E., Heuser, J.E., Kerkhoff, E., and Dyche Mullins, R. (2005). Drosophila Spire is an actin nucleation factor. Nature 433, 382–388. 10.1038/nature03241.

R Core Team (2021). R: A Language and Environment for Statistical Computing.

Ray, R.P., Matamoro-Vidal, A., Ribeiro, P.S., Tapon, N., Houle, D., Salazar-Ciudad, I., and Thompson, B.J. (2015). Patterned Anchorage to the Apical Extracellular Matrix Defines Tissue Shape in the Developing Appendages of Drosophila. Dev. Cell 34, 310–322. 10.1016/j.devcel.2015.06.019.

Reed, R.D. (2004). Evidence for Notch-mediated lateral inhibition in organizing butterfly wing scales. Dev. Genes Evol. 214, 43–46. 10.1007/s00427-003-0366-0.

Schweisguth, F. (2015). Asymmetric cell division in the Drosophila bristle lineage: From the polarization of sensory organ precursor cells to Notch-mediated binary fate decision. Wiley Interdiscip. Rev. Dev. Biol. 4, 299–309. 10.1002/wdev.175.

Sekine, Y., Takagahara, S., Hatanaka, R., Watanabe, T., Oguchi, H., Noguchi, T., Naguro, I., Kobayashi, K., Tsunoda, M., Funatsu, T., et al. (2011). p38 MAPKs regulate the expression of genes in the dopamine synthesis pathway through phosphorylation of NR4A nuclear receptors. J. Cell Sci. 124, 3006–3016. 10.1242/jcs.085902.

Stuart, T., Butler, A., Hoffman, P., Hafemeister, C., Papalexi, E., Mauck, W.M., Hao, Y., Stoeckius, M., Smibert, P., and Satija, R. (2019). Comprehensive Integration of Single-Cell Data. Cell 177, 1888–1902.e21. 10.1016/j.cell.2019.05.031.

Tian, S., and Monteiro, A. (2022). A Transcriptomic Atlas Underlying Developmental Plasticity of Seasonal Forms of Bicyclus anynana Butterflies. Mol. Biol. Evol. 39, msac126. 10.1093/molbev/msac126.

Tian, L., Su, S., Dong, X., Amann-Zalcenstein, D., Biben, C., Seidi, A., Hilton, D.J., Naik, S.H., and Ritchie, M.E. (2018). scPipe: A flexible R/Bioconductor preprocessing pipeline for single- cell RNA-sequencing data. PLoS Comput. Biol. 14, 1–15. 10.1371/journal.pcbi.1006361.

Vanhoutteghem, A., Maciejewski-Duval, A., Bouche, C., Delhomme, B., Hervé, F., Daubigney, F., Soubigou, G., Araki, M., Araki, K., Yamamura, K., et al. (2009). Basonuclin 2 has a function in the multiplication of embryonic craniofacial mesenchymal cells and is orthologous to disco proteins. Proc. Natl. Acad. Sci. 106, 14432–14437. 10.1073/pnas.0905840106.

Xie, B., Charlton-Perkins, M., McDonald, E., Gebelein, B., and Cook, T. (2007). Senseless functions as a molecular switch for color photoreceptor differentiation in Drosophila. Development 134, 4243–4253. 10.1242/dev.012781.

Yao, L., Wang, S., Westholm, J.O., Dai, Q., Matsuda, R., Hosono, C., Bray, S., Lai, E.C., and Samakovlis, C. (2017). Genome-wide identification of Grainy head targets in Drosophila reveals regulatory interactions with the POU domain transcription factor Vvl. 3145–3155. 10.1242/dev.143297.

Yoshida, A., and Emoto, J. (2011). Variations in the arrangement of sensory bristles along butterfly wing margins. Zoolog. Sci. 28, 430–437. 10.2108/zsj.28.430.

Yoshida, A., Noda, A., and Emoto, J. (2001). Bristle distribution along the wing margin of the small white cabbage butterfly (Lepidoptera: Pieridae). Ann. Entomol. Soc. Am. 94, 467–470. 10.1603/0013-8746(2001)094[0467:BDATWM]2.0.CO;2.

Yu, G., Wang, L.-G., Han, Y., and He, Q.-Y. (2012). clusterProfiler: an R Package for Comparing Biological Themes Among Gene Clusters. Omi. A J. Integr. Biol. 16, 284–287. 10.1089/omi.2011.0118.

Zeng, H., and Sanes, J.R. (2017). Neuronal cell-type classification: Challenges, opportunities and the path forward. Nat. Rev. Neurosci. 18, 530–546. 10.1038/nrn.2017.85.

Zhang, Y.-N., Zhu, X.-Y., Fang, L.-P., He, P., Wang, Z.-Q., Chen, G., Sun, L., Ye, Z.-F., Deng, D.-G., and Li, J.-B. (2015). Identification and Expression Profiles of Sex Pheromone Biosynthesis and Transport Related Genes in Spodoptera litura. PLoS One 10, 1–22. 10.1371/journal.pone.0140019.

Zhou, Q., Zhang, T., Xu, W., Yu, L., Yi, Y., and Zhang, Z. (2008). Analysis of four achaete- scute homologs in Bombyx mori reveals new viewpoints of the evolution and functions of this gene family. BMC Genet. 9, 1–13. 10.1186/1471-2156-9-24.

Zhou, Q., Yu, L., Shen, X., Li, Y., Xu, W., Yi, Y., and Zhang, Z. (2009). Homology of dipteran bristles and lepidopteran scales: Requirement for the Bombyx mori achaete-scute homologue ASH2. Genetics 183, 619–627. 10.1534/genetics.109.102848.

