## Supplementary Materials for "The molecular basis of macrochaete diversification highlighted by a single-cell atlas of *Bicyclus anynana* butterfly pupal forewings"

**This file includes:**

Supplementary Text
References for SI
Supplementary Figures S1-S16
Supplementary Tables S1-S5

### Supplementary Text

Macrochaete development has been well studied in the fruit fly *Drosophila melanogaster*. In flies, each macrochaete consists of four different cell types – the external socket and bristle cells and the internal sheath and neuron cells – that arise from a single sensory organ precursor (SOP) cell (Hartenstein and Posakony, 1990). Initially, this SOP is selected from a proneural cluster of cells by the action of various proneural genes of the Achaete-scute complex (AS-C) (Artavanis-Tsakonas and Simpson, 1991; Furman and Bukharina, 2008). The SOP promotes an epidermal cell fate in neighboring cells by repressing AS-C genes through a Notch-mediated lateral inhibition process (Artavanis-Tsakonas and Simpson, 1991; Hartenstein and Posakony, 1990; Simpson, 1990). The SOP then undergoes two rounds of asymmetric cell division to produce four daughter cells (Hartenstein and Posakony, 1990; Schweisguth, 2015) (Fig 1A). The first division leads to two daughter cells, pIIa and pIIb, where pIIa divides to form the future socket and bristle cells, and pIIb divides into the neuron and sheath cells. At each cell division, binary cell fate choices are regulated through Notch signaling, initiated by the asymmetric distribution of cell fate determinants like *numb* and *neuralized (neur)* (Artavanis-Tsakonas and Simpson, 1991; Le Borgne and Schweisguth, 2003; Hartenstein and Posakony, 1990; Rhyu et al., 1994; Uemura et al., 1989). In the first division, *numb* and *neur* are asymmetrically segregated to the pIIb cell which acts as the signal sending cell, activating Notch in the pIIa cell (Fig 1A). In the second set of divisions, the socket and sheath cells exhibit active Notch signaling (higher levels of Notch and its downstream targets), initiated by the bristle and neuron cell, respectively (Le Borgne and Schweisguth, 2003; Rhyu et al., 1994) (Fig 1A).

The roles of various genes in driving cell fate regulation and cell differentiation in the sensory organ lineage in *Drosophila*, after SOP specification, have been well-characterized. For example, *cut* is expressed in the SOP cell of all external sensory organs and its progeny but is not expressed in the internal chordotonal sensory organs (Blochlinger et al., 1988, 1990). Similarly, *couch potato* (*cpo*) is expressed in all cells of the peripheral nervous system and requires the expression of *achaete-scute* and *daughterless* (Bellen et al., 1992). The transcription factor *tramtrack* (*ttk*) is expressed in all non-neuronal cells during cell fate specification (Guo et al., 1995). Members of the Notch signaling pathway such as *Notch, Delta and Enhancer of split* [*E(spl)*] play key roles in cell fate determination (Artavanis-Tsakonas and Simpson, 1991). *scabrous* (*sca*) is a secreted protein that is highly expressed in the sensory organ precursors on the wing imaginal discs of *Drosophila*. sca mutants show an increased bristle density with a disordered bristle organization and sometimes a double bristle phenotype due to incorrect cell fate specification (Mlodzik et al., 1990; Renaud and Simpson, 2001). A homolog of the vertebrate *Pax2* gene, *D-Pax2*, corresponding to the shaven locus in *Drosophila*, is initially expressed in the SOP cell and its daughters pIIa and pIIb but is later highly expressed and restricted to the shaft and sheath cells only (Kavaler et al., 1999). *elav* and *prospero* are markers of the pIIb progeny cells i.e., the neuron and sheath cells (Bier et al., 1988; Manning and Doe, 1999).

<u>References</u>

Artavanis-Tsakonas, S., and Simpson, P. (1991). Choosing a cell fate: a view from the Notch locus. Trends Genet. *7*, 403–408. https://doi.org/10.1016/0168-9525(91)90264-q.

Bellen, H.J., Kooyer, S., D'Evelyn, D., and Pearlman, J. (1992). The Drosophila couch potato protein is expressed in nuclei of peripheral neuronal precursors and shows homology to RNA-binding proteins. Genes Dev. *6*, 2125–2136. https://doi.org/10.1101/gad.6.11.2125.

Bier, E., Ackerman, L., Barbel, S., Jan, L., and Jan, Y.N. (1988). Identification and characterization of a neuron-specific nuclear antigen in Drosophila. Science *240*, 913–916. https://doi.org/10.1126/science.3129785.

Blochlinger, K., Bodmer, R., Jack, J., Jan, L.Y., and Jan, Y.N. (1988). Primary structure and expression of a product from cut, a locus involved in specifying sensory organ identity in Drosophila. Nature *333*, 629–635. https://doi.org/10.1038/333629a0.

Blochlinger, K., Bodmer, R., Jan, L.Y., and Jan, Y.N. (1990). Patterns of expression of cut, a protein required for external sensory organ development in wild-type and cut mutant Drosophila embryos. Genes Dev. *4*, 1322–1331. https://doi.org/10.1101/gad.4.8.1322.

Le Borgne, R., and Schweisguth, F. (2003). Unequal segregation of neuralized biases Notch activation during asymmetric cell division. Dev. Cell *5*, 139–148. https://doi.org/10.1016/S1534-5807(03)00187-4.

Furman, D., and Bukharina, T. (2008). How Drosophila melanogaster Forms its Mechanoreceptors. Curr. Genomics *9*, 312–323. https://doi.org/10.2174/138920208785133271.

Guo, M., Bier, E., Jan, L.Y., and Jan, Y.N. (1995). tramtrack acts downstream of numb to specify distinct daughter cell fates during asymmetric cell divisions in the drosophila PNS. Neuron *14*, 913–925. https://doi.org/https://doi.org/10.1016/0896-6273(95)90330-5.

Hartenstein, V., and Posakony, J.W. (1990). A dual function of the Notch gene in Drosophila sensillum development. Dev. Biol. *142*, 13–30. https://doi.org/10.1016/0012-1606(90)90147-b.

Kavaler, J., Fu, W., Duan, H., Noll, M., and Posakony, J.W. (1999). An essential role for the Drosophila Pax2 homolog in the differentiation of adult sensory organs. Development *126*, 2261–2272. https://doi.org/10.1242/dev.126.10.2261.

Manning, L., and Doe, C.Q. (1999). Prospero distinguishes sibling cell fate without asymmetric localization in the Drosophila adult external sense organ lineage. Development *126*, 2063–2071. https://doi.org/10.1242/dev.126.10.2063.

Mlodzik, M., Baker, N.E., and Rubin, G.M. (1990). Isolation and expression of scabrous , a gene regulating neurogenesis in Drosophila. 1848–1861. .

Renaud, O., and Simpson, P. (2001). scabrous Modifies Epithelial Cell Adhesion and Extends the Range of Lateral Signalling during Development of the Spaced Bristle Pattern in Drosophila. Dev. Biol. *240*, 361–376. https://doi.org/https://doi.org/10.1006/dbio.2001.0482.

Rhyu, M.S., Jan, L.Y., and Jan, Y.N. (1994). Asymmetric distribution of numb protein during division of the sensory organ precursor cell confers distinct fates to daughter cells. Cell *76*, 477–491. https://doi.org/10.1016/0092-8674(94)90112-0.

Schweisguth, F. (2015). Asymmetric cell division in the Drosophila bristle lineage: From the polarization of sensory organ precursor cells to Notch-mediated binary fate decision. Wiley Interdiscip. Rev. Dev. Biol. *4*, 299–309. https://doi.org/10.1002/wdev.175.

Simpson, P. (1990). Lateral inhibition and the development of the sensory bristles of the adult peripheral nervous system of Drosophila. Development *109*, 509–519. https://doi.org/10.1242/dev.109.3.509.

Uemura, T., Shepherd, S., Ackerman, L., Jan, L.Y., and Jan, Y.N. (1989). numb, a gene required in determination of cell fate during sensory organ formation in Drosophila embryos. Cell *58*, 349–360. https://doi.org/https://doi.org/10.1016/0092-8674(89)90849-0.

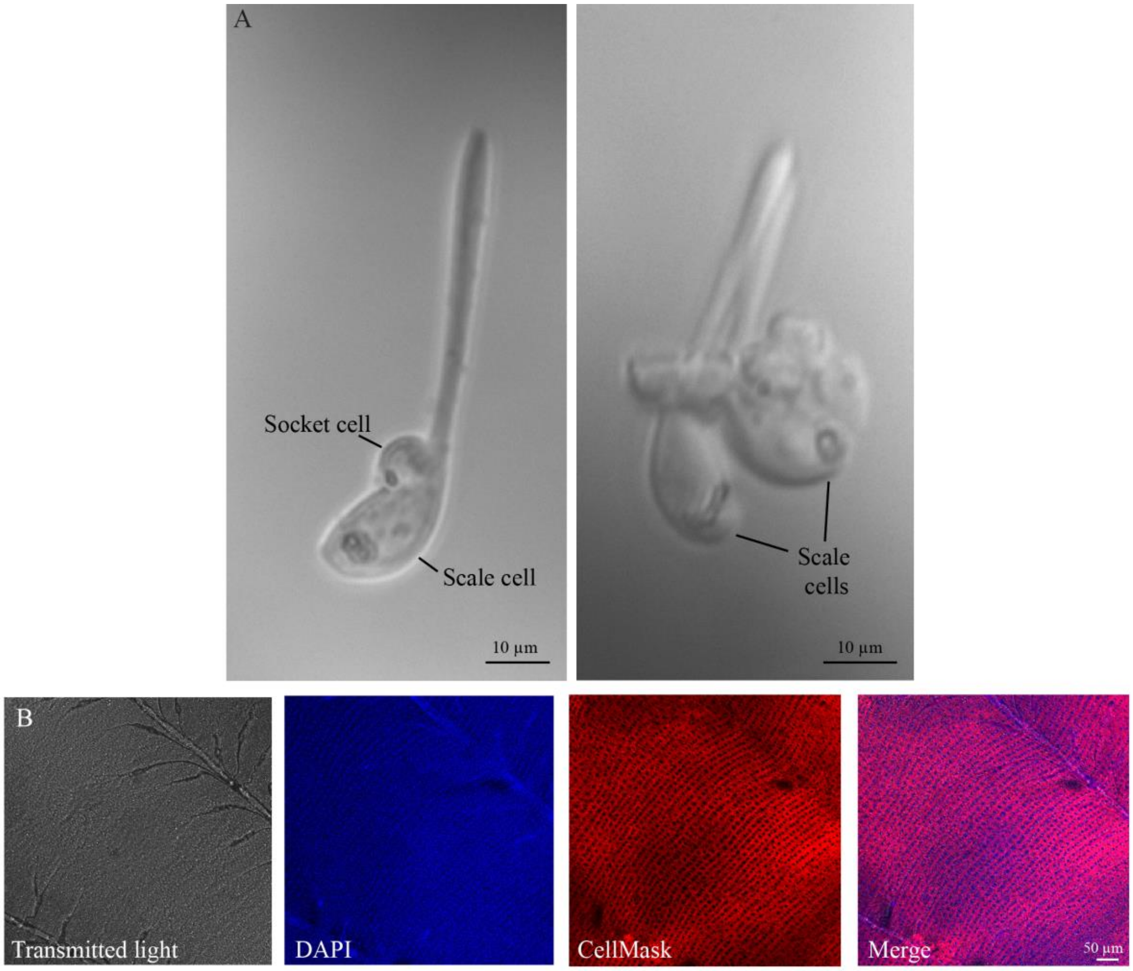


**Figure S1**: **Scales and scale precursor cells in *Bicyclus anynana***. (A) Isolated scale and socket cells imaged from 48-72 hours dissociated forewings of *B. anynana*. Extensions of the scale cells that become the future adult scales are visible. Scale bars are 10 µm. (B) A 24-hour pupal wing disc of *B. anynana* stained with DAPI to image the nucleus and CellMask Plasma Membrane stain to image the plasma membrane. Neat rows of scale precursor cells >6 µm in diameter are visible. Scale bar is 50 µm.

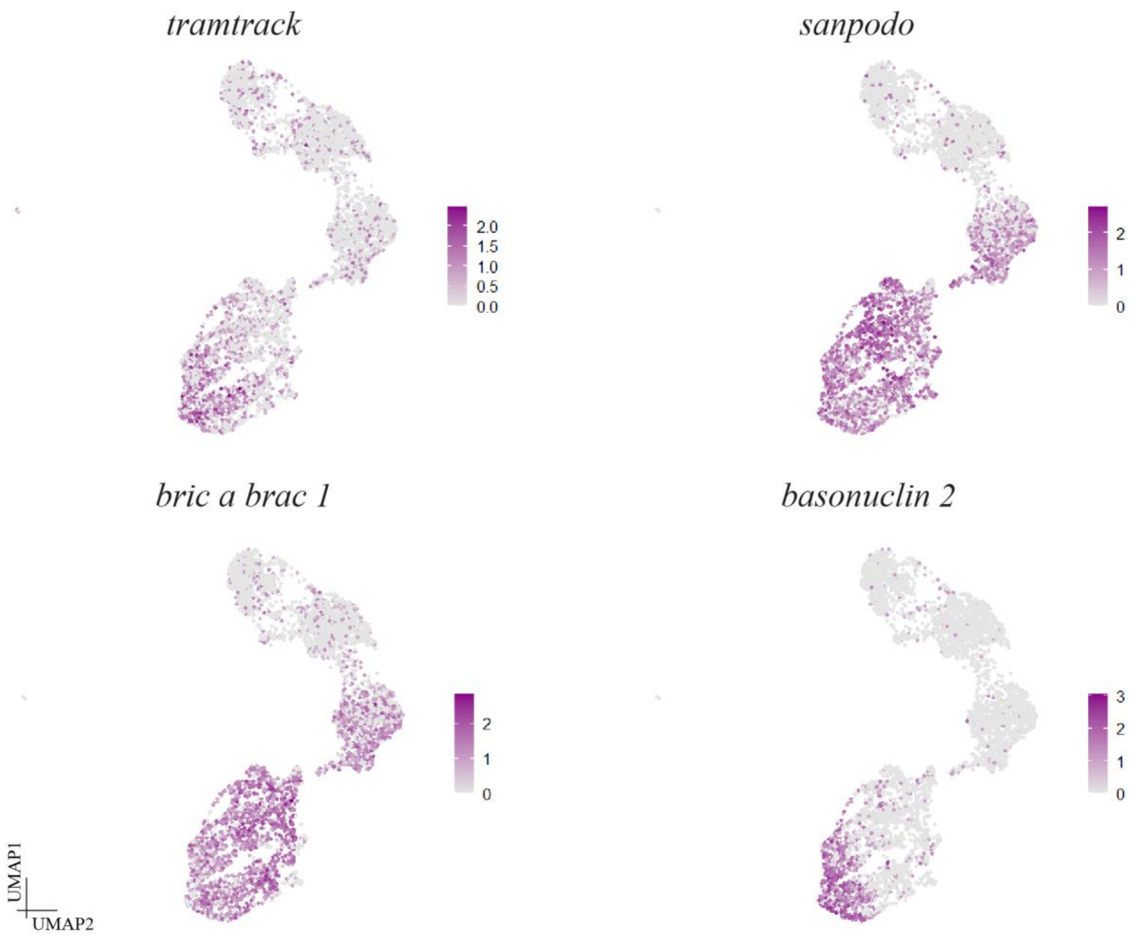


**Figure S2**: **UMAP expression plots of genes highly expressed in the scale cell clusters.** *tramtrack* expression is similar to *cut* and *scabrous*. *sanpodo* and *bric a brac 1* are broadly expressed in the scale cell clusters, similar to *cpo*, while *basonuclin 2* is highly expressed in a subset of the scale cell class.

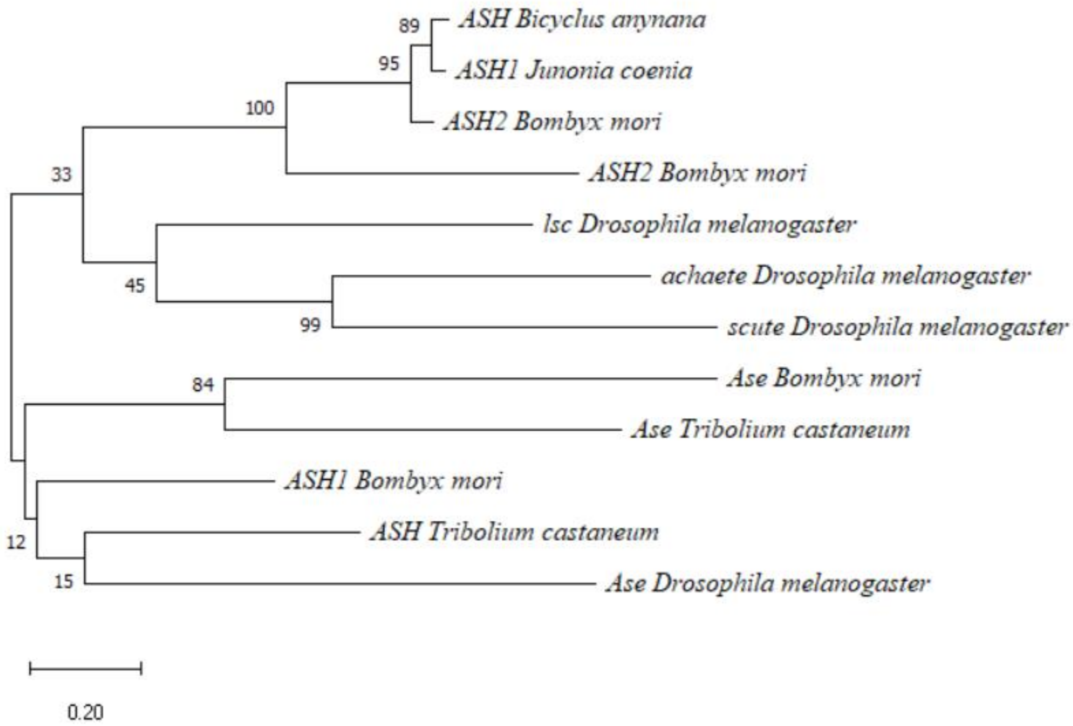


**Figure S3**: **Phylogenetic tree based on the protein sequences of the *achaete-scute homologs***. The evolutionary relatedness of the two butterfly *achaete-scute homologs* to other Lepidopteran, Dipteran and Coleopteran AS-C genes was identified using a Maximum Likelihood tree in MEGA X. The percentage of trees in which the associated taxa clustered together is shown next to the branches. The tree is drawn to scale, with branch lengths measured in the number of substitutions per site. Both butterfly *ASH* genes cluster with *Bombyx mori ASH2*.

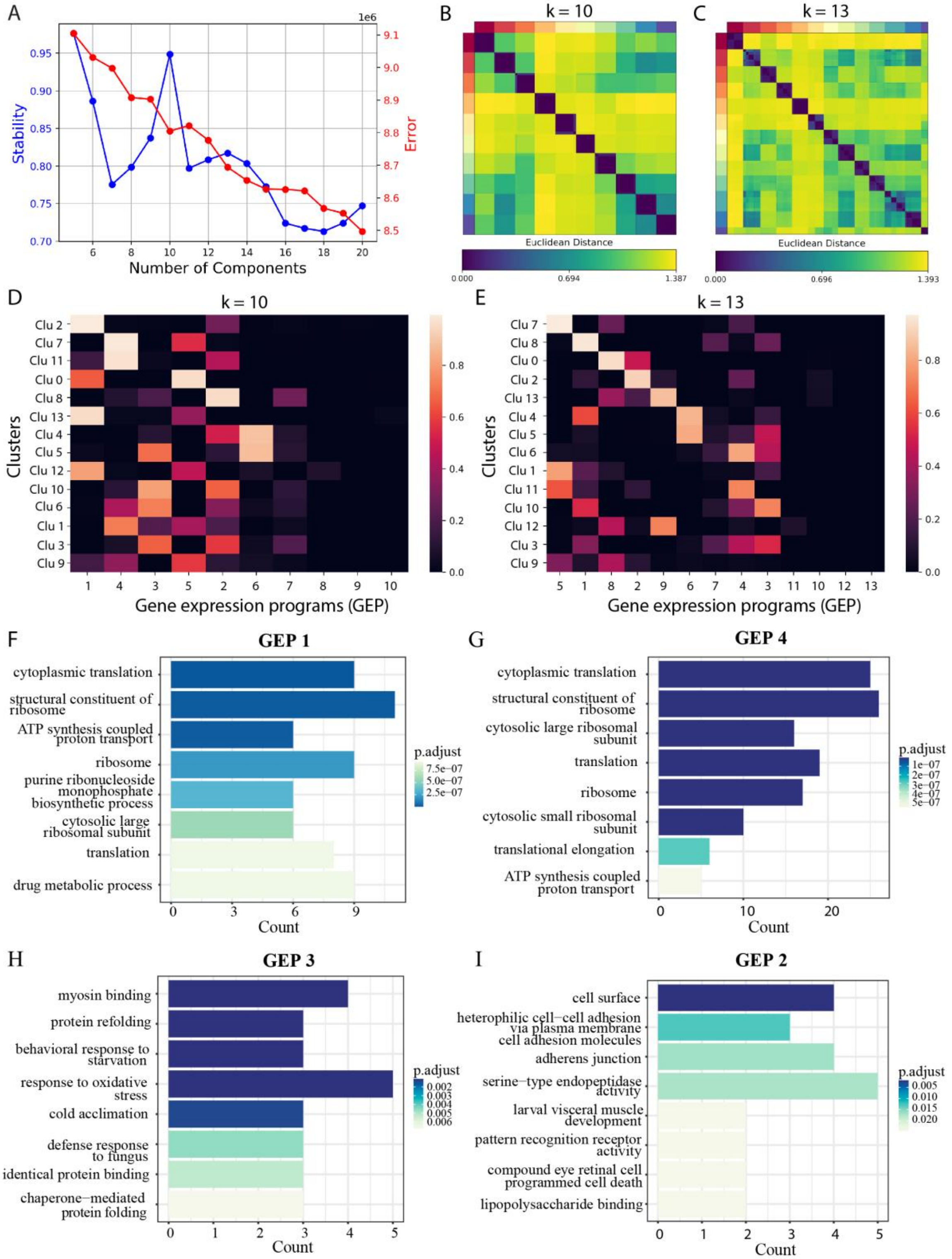


**Figure S4**: **K selection in cNMF**. (A) Plot of the stability (blue line) and the Frobenius reconstruction error as a function of the number of cNMF components (K) using a dataset with 2000 over-dispersed genes. (B, C) Consensus estimate plots for K = 10 and 13 with outlier filtering (threshold of 0.18) showing good concordance between the 200 cNMF replicates. (D, E) Usage maps of the contributions of 10 vs 13 GEPs to the 14 different cell types. K=10 provides a poorer resolution across cell types as compared to K=13. (F-I) Gene enrichment plots of the top 60 genes within (F) GEP 1, (G) GEP 4, (H) GEP 3 and (I) GEP 2.

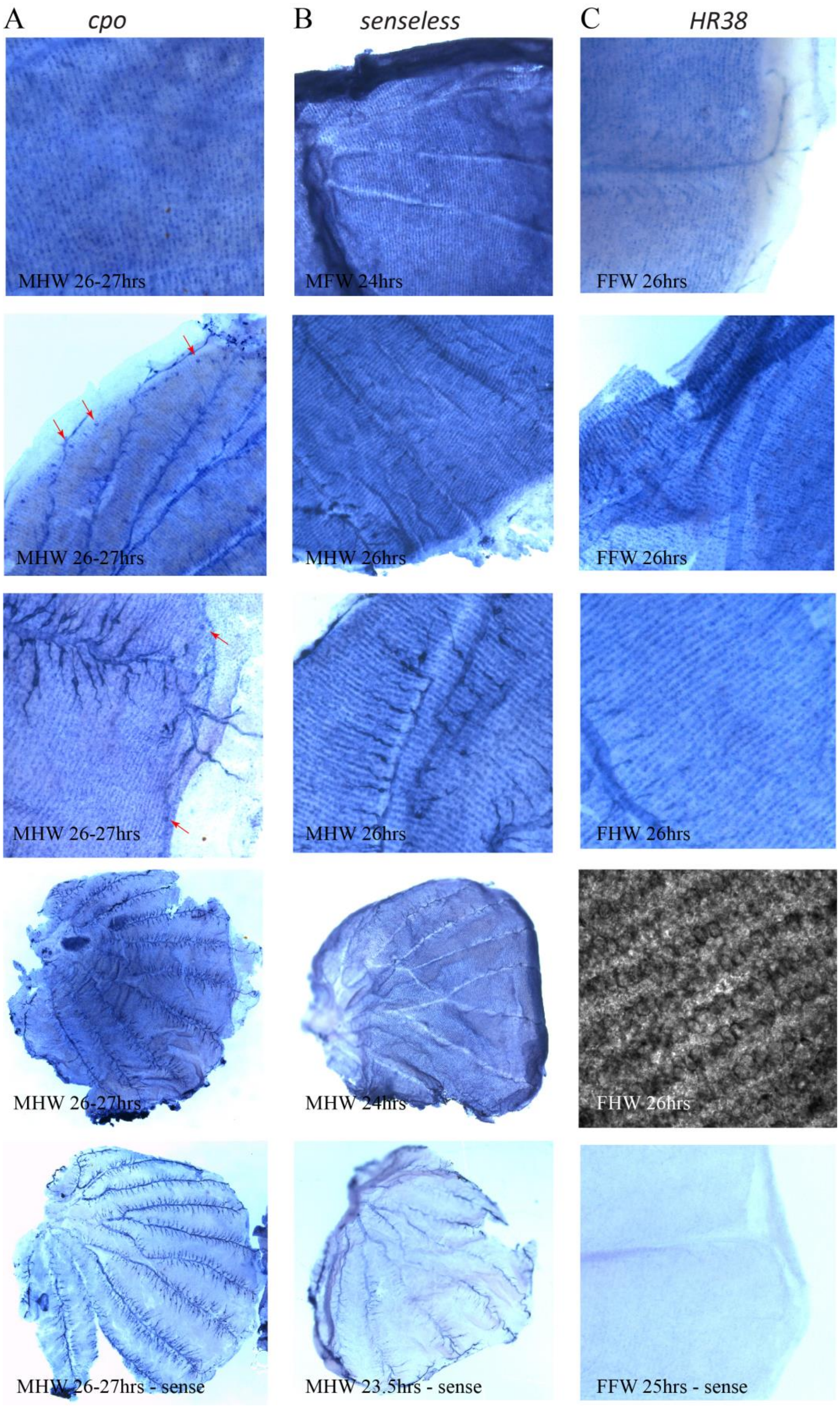
A cpo
B senseless
C HR38
MHW 26-27hrs
MFW 24hrs
FFW 26hrs
MHW 26-27hrs
MHW 26hrs
FFW 26hrs
MHW 26-27hrs
MHW 26hrs
FHW 26hrs
MHW 26-27hrs
MHW 24hrs
FHW 26hrs
MHW 26-27hrs - sense
MHW 23.5hrs - sense
FFW 25hrs - sense

**Figure S5**: ***in situ* hybridization of 25-28 hour developing pupal wing discs of *Bicyclus anynana* using probes against *cpo*, *senseless* and *HR38***. Control stains with a sense probe for each gene are shown in the last row. (A) *cpo* is expressed in the cells of the scale cell lineage across the wing. It is additionally very highly expressed in the hair-pencil cells of the male hindwing (row 4) and in the wing margin mechanosensory cells (red arrows). (B) Expression of *senseless* is seen in neat rows across the pupal wing. (C) Expression of *HR38* is also seen in neat rows on the wing. The image in row 4 was taken under the transmitted light setting of a confocal microscope. For all three genes, using *in situ* hybridization we couldn't determine if these cells were the pIIa cells or the daughter cells but based on the size of the cells, they are potentially the daughter scale cells. MFW/FFW: Male/Female forewing; MHW/FHW: Male/Female hindwing.

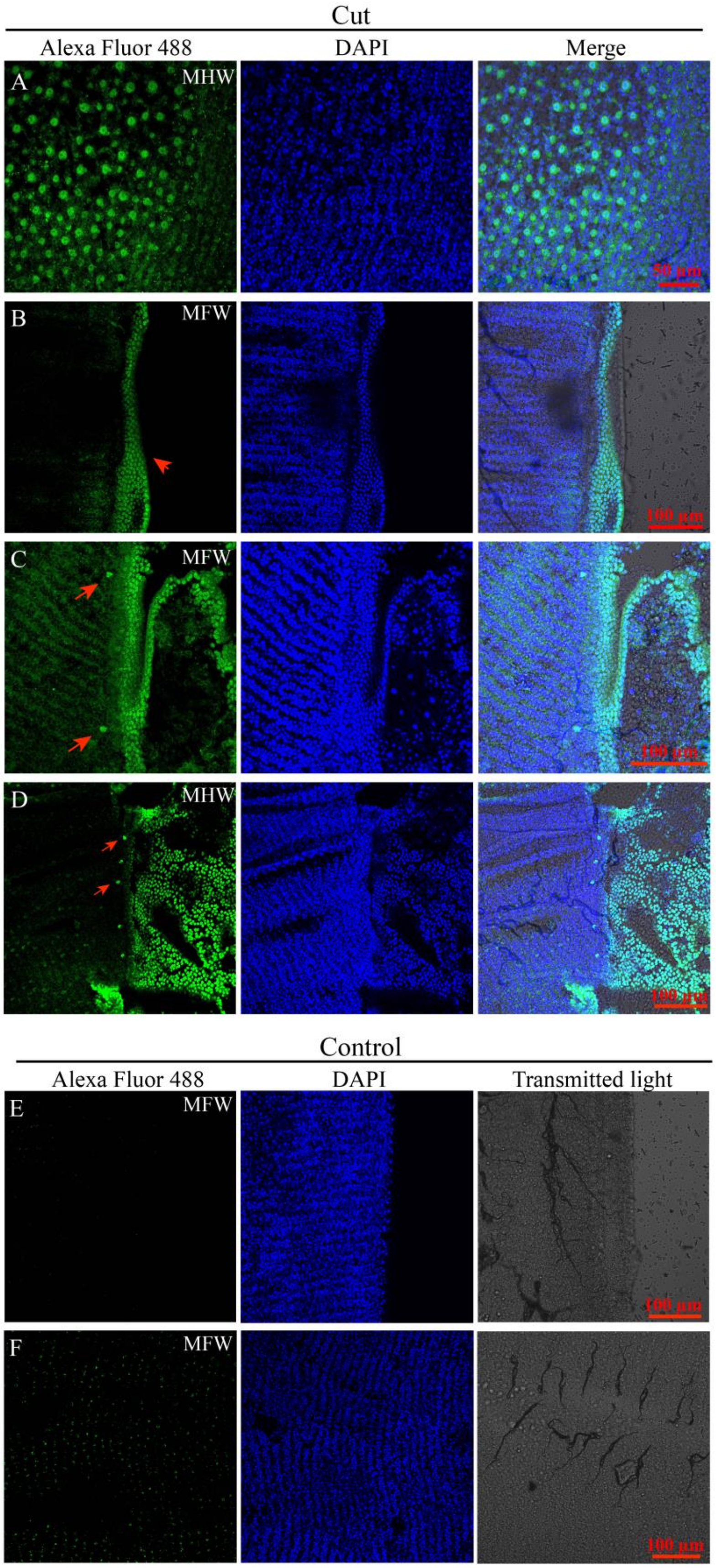
Cut
Alexa Fluor 488
DAPI
Merge
A
MHW
50 µm
B
MFW
100 µm
C
MFW
100 µm
D
MHW
100 µm
Control
Alexa Fluor 488
DAPI
Transmitted light
E
MFW
100 µm
F
MFW
100 µm

**Figure S6**: **Cut protein expression in 25-28 hour developing pupal wing discs of *Bicyclus anynana***. Cut expression is seen in the (A) cytoplasm of the long, hair-like scales on the hindwing (B) in the cells of the peripheral tissue close to the wing margin, which will undergo apoptosis (red arrow) and (C, D) in the marginal mechanosensory bristles (red arrows). (E, F) Control stains with only secondary antibody. MFW: Male forewing; MHW: Male hindwing.

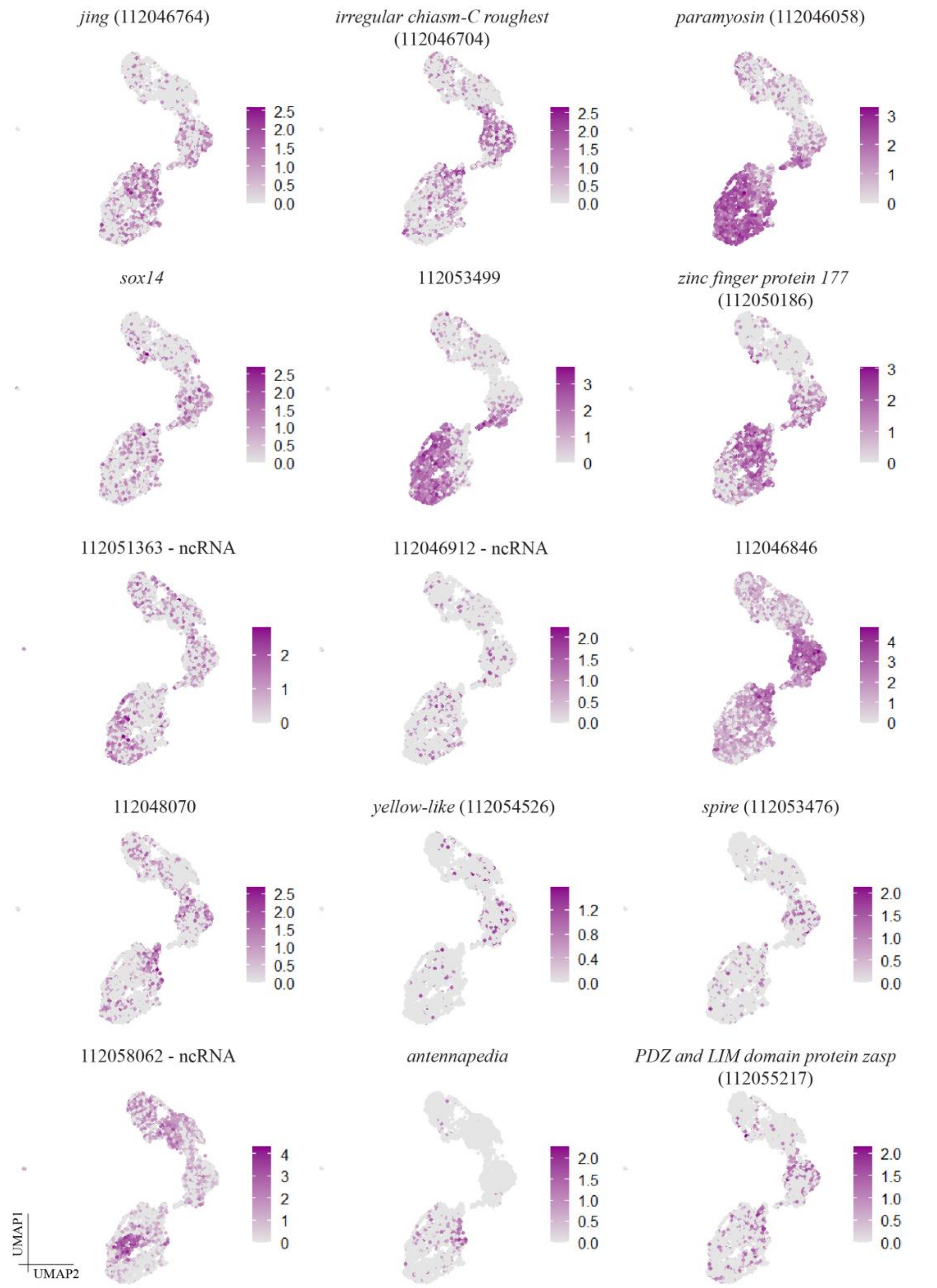


**Figure S7**: **UMAP expression plots of some of the highly expressed genes in the scale cell clusters.** NCBI GeneIDs are provided in parenthesis. Labels that begin with 1120 correspond to the NCBI GeneIDs of uncharacterized genes and ncRNA refers to non-coding RNA.

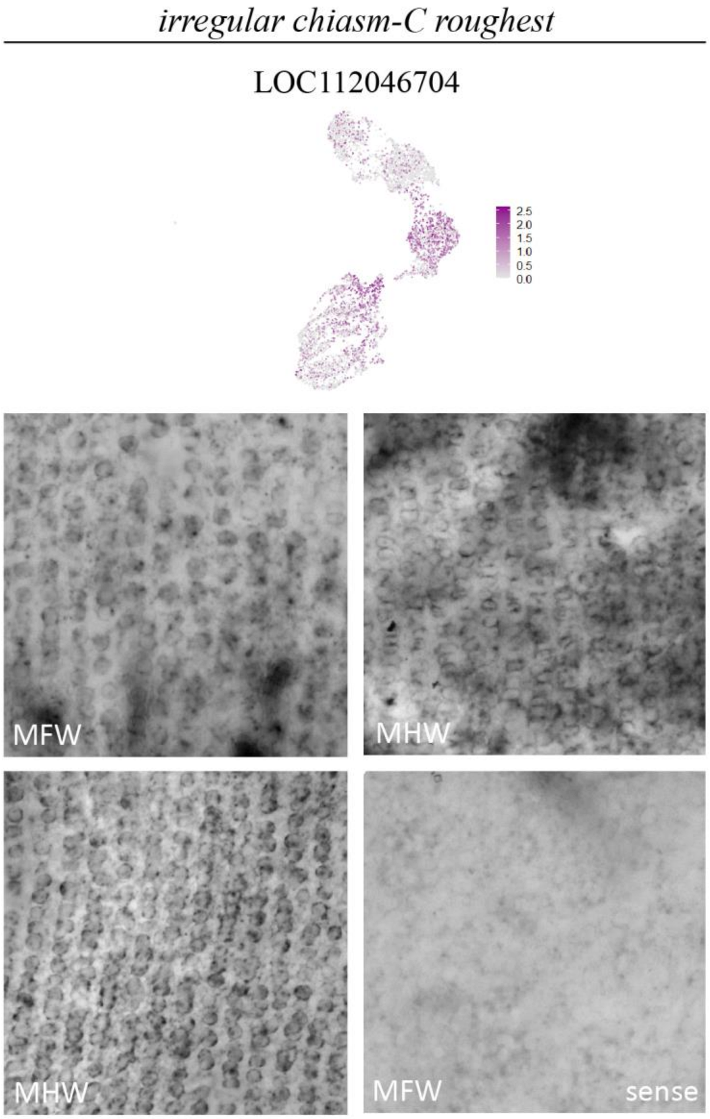


**Figure S8**: ***irregular chiasm C-roughest (rst)* mRNA is expressed in the scale cells across the pupal wing.** One *rst* gene (NCBI Gene ID - LOC112046704) is expressed in the scale cells across 25-28 hour pupal wings. Bottom right panel is a sense stain. MFW: Male forewing; MHW: Male hindwing.

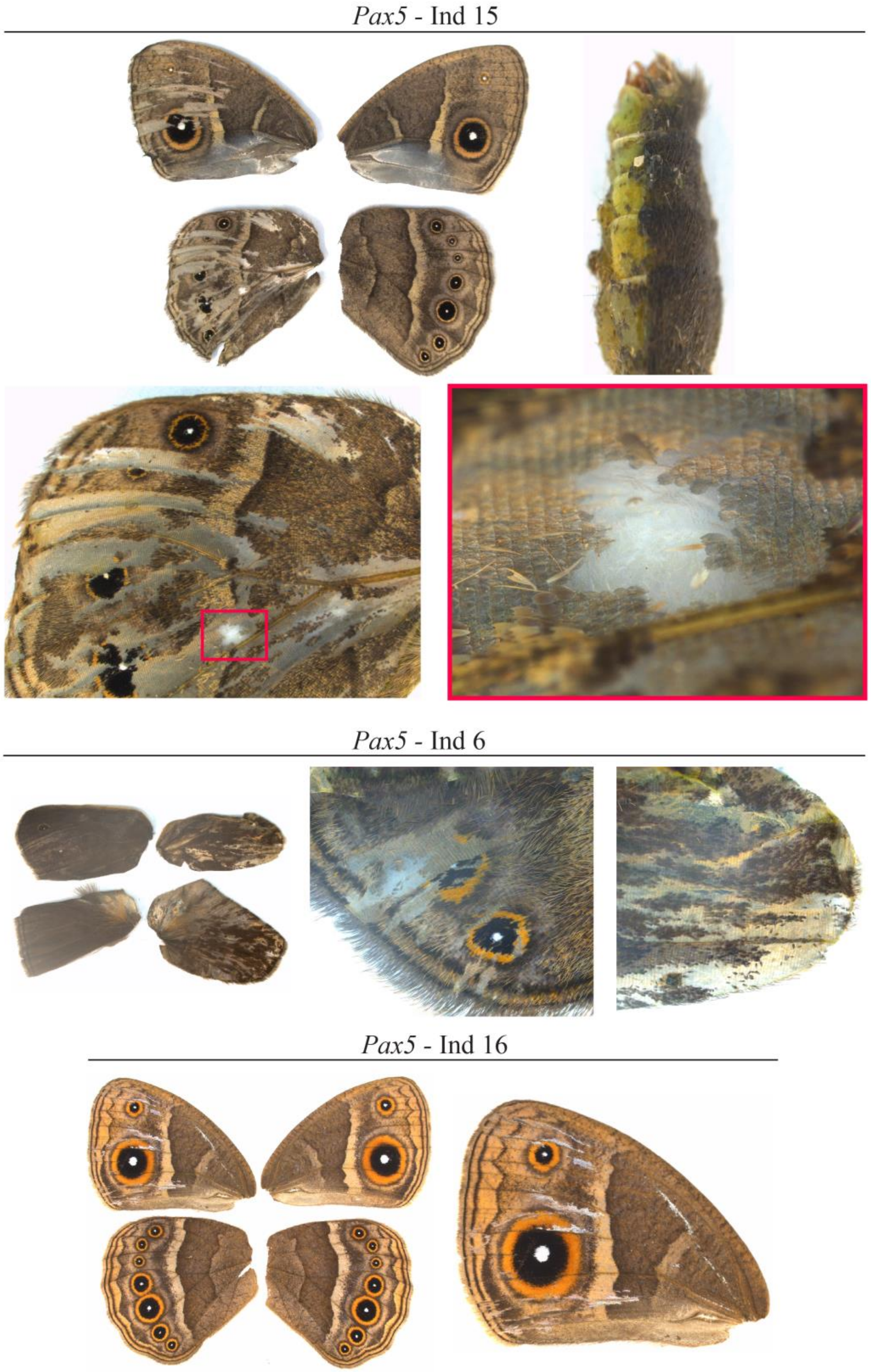


**Figure S9: *Pax5* crispants of *B. anynana*.** Representative images of various *Pax2* crispants. Colored, boxed area is magnified to show loss of scales.

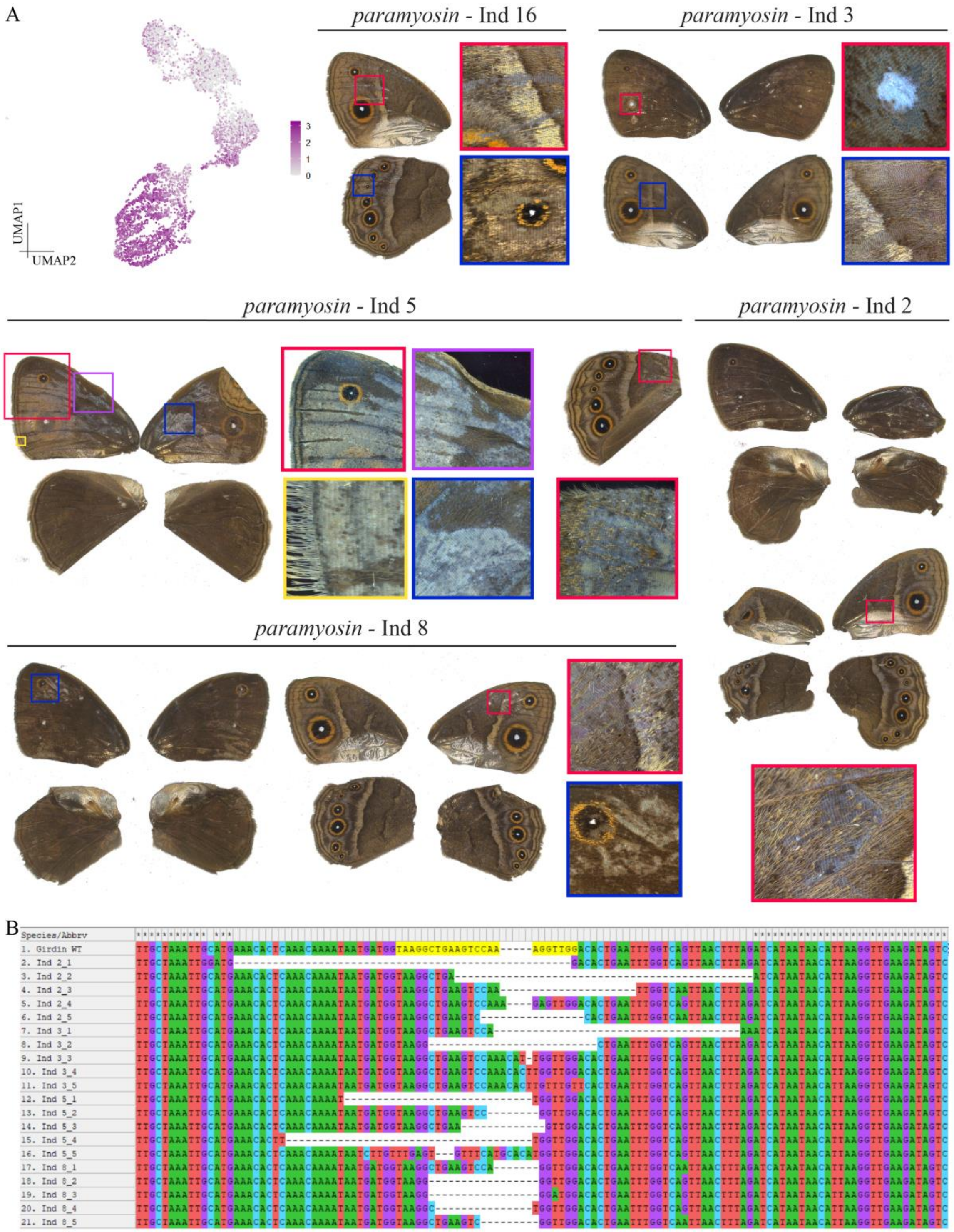


**Figure S10**: ***paramyosin* is important for scale cell development**. (A) UMAP expression plot of *paramyosin*. It is highly expressed in cells of the scale cell lineage. *paramyosin* crispants of *B. anynana* exhibit loss of scales on both dorsal and ventral surfaces. Cover scales are lost more often than ground scales. (B) Mutations in the targeted region of *paramyosin* across various crispant individuals.

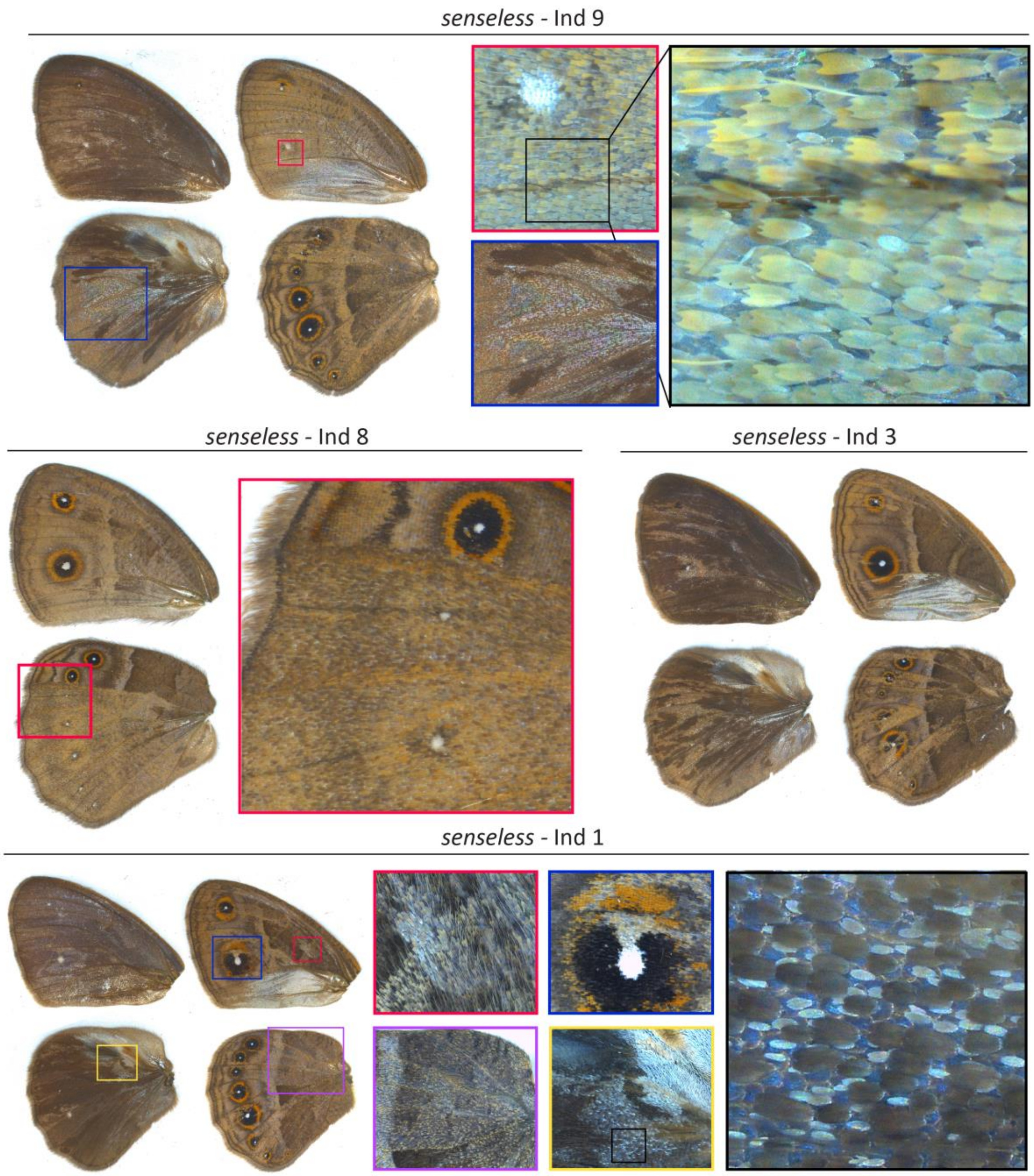


**Figure S11: *senseless* crispants of *B. anynana*.** Representative images of various *senseless* crispants. Colored, boxed areas are magnified to show loss or transformations of scales.

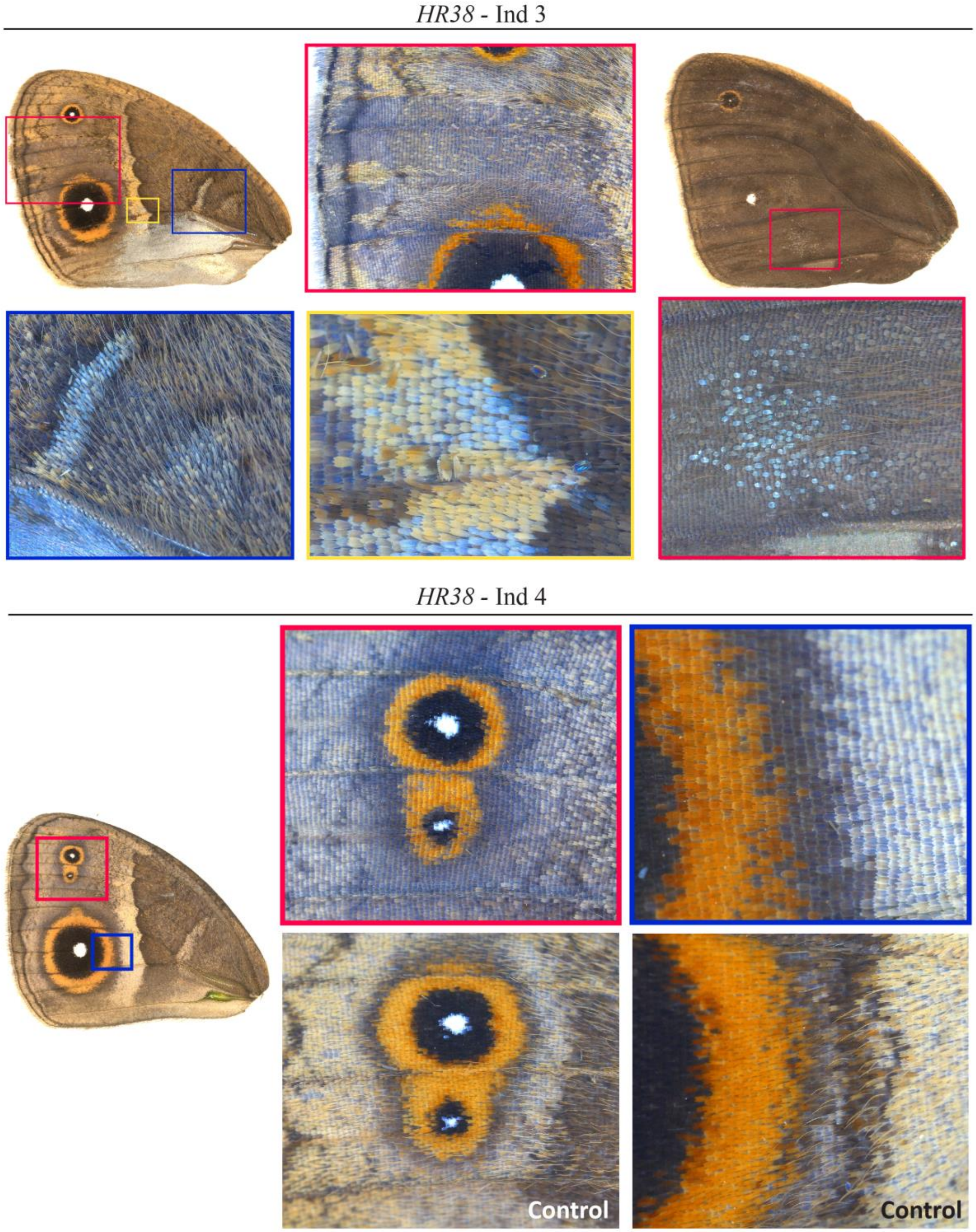


**Figure S12: *HR38* crispants of *B. anynana*.** Representative images of *HR38* crispants. Colored, boxed areas are magnified to show loss or transformations of scale types.

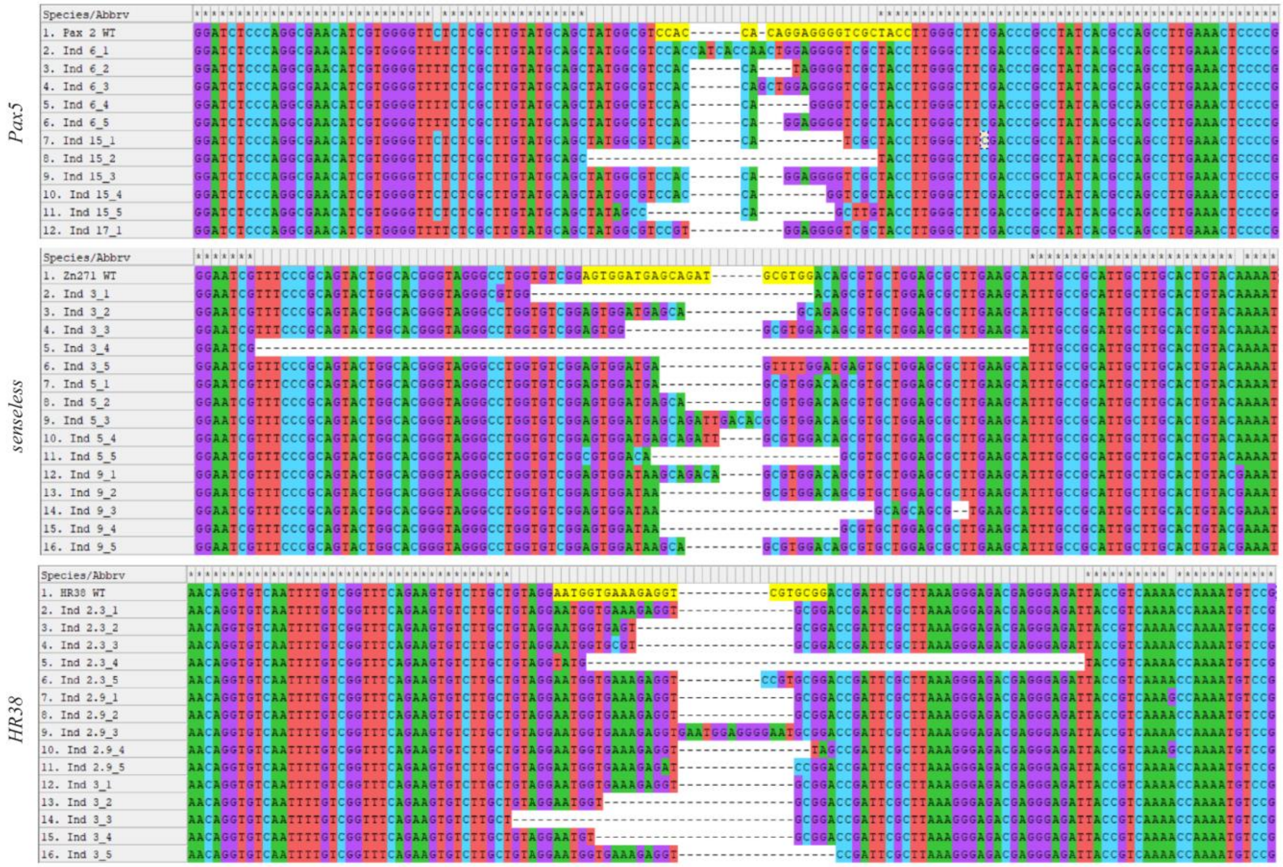


**Figure S13: Genotyping crispant individuals.** Mutations in the targeted regions of *Pax5*, *senseless* and *HR38* in various crispant individuals.

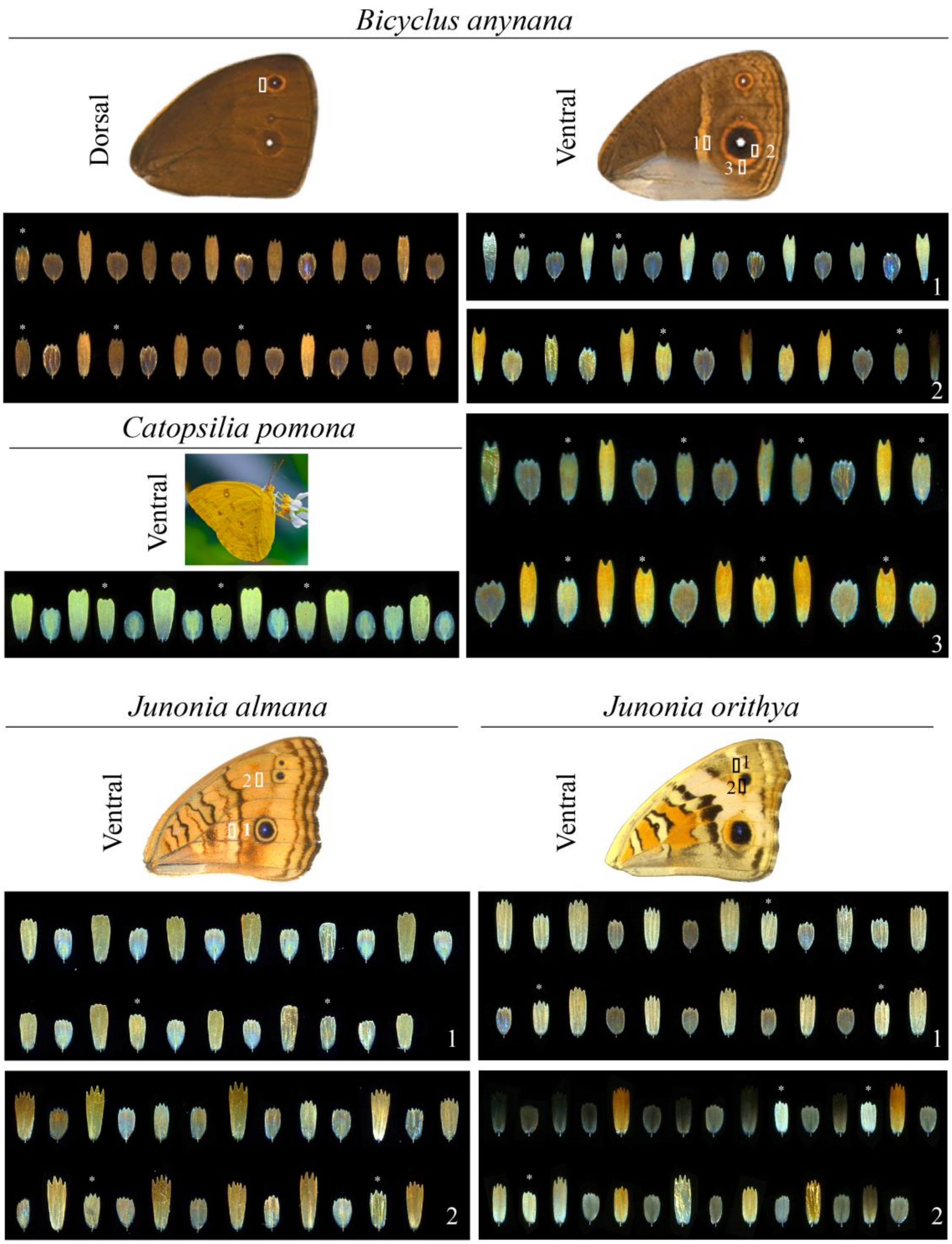


**Figure S14: Scale organization in various butterfly species.** Scale organization in various parts of the dorsal or ventral wings of *Bicyclus anynana*, *Catopsilia pomona*, *Junonia almana* and *Junonia orithya* butterflies. Apart from the cover and ground scales, an intermediate scale type is often visible (marked with white stars). Location on the wings from which the scales were picked are marked with numbers and boxes. *Junonia almana* forewing image taken by Jocelyn Wee and *Catopsilia pomona* image captured by Victoria Long.

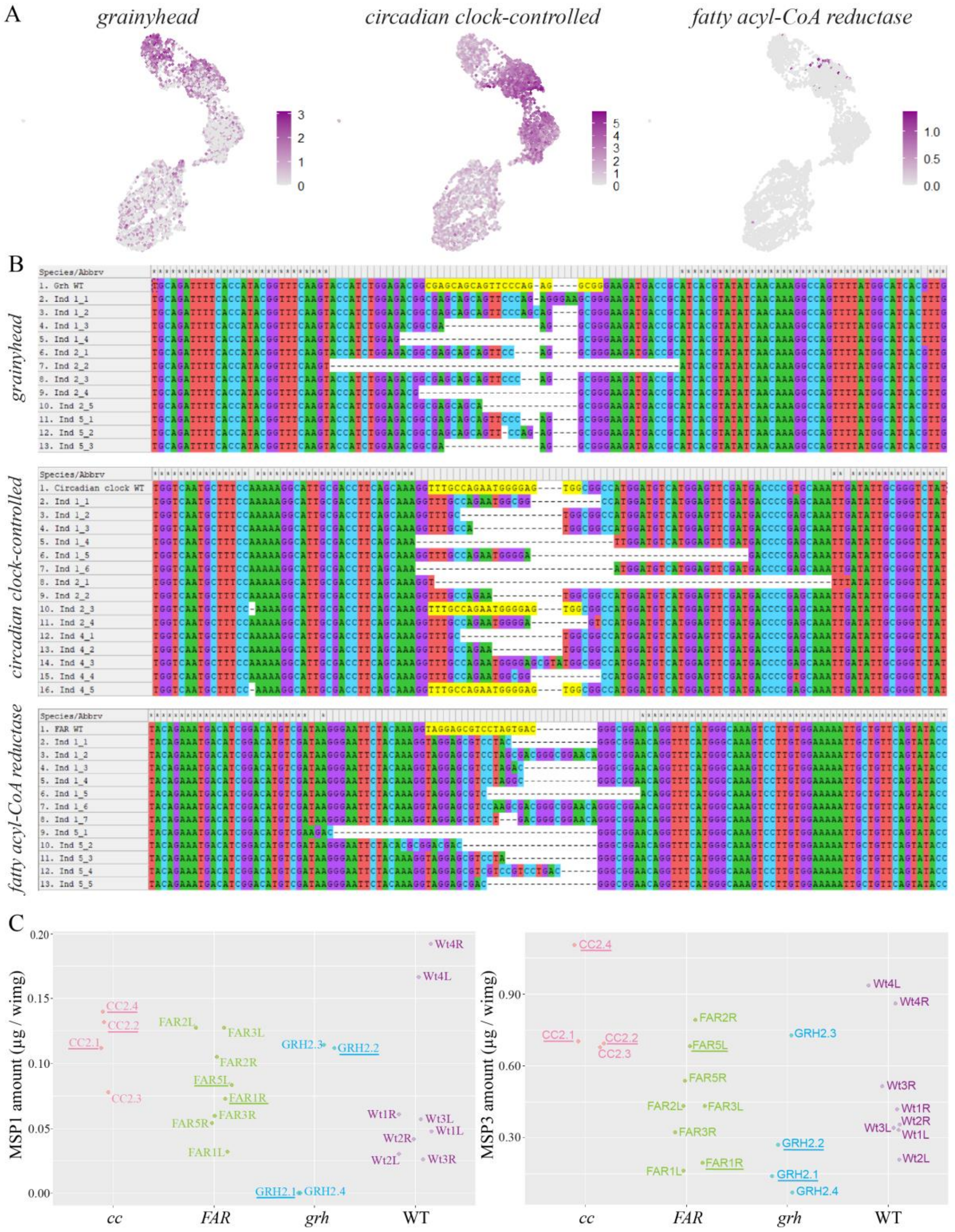


**Figure S15: *grainyhead, circadian clock-controlled* and *fatty acyl-CoA reductase* expression and pheromone composition in *B. anynana* crispants.** (A) UMAP expression plots of *grainyhead, circadian clock-controlled and fatty acyl-CoA reductase* transcripts. (B) Mutations in the targeted regions of the three genes in various crispant individuals. (C) MSP1 and MSP3 amounts extracted from the wings of 3-day old males from a batch of CRISPR-Cas9 + respective guide RNA injected individuals. Each data point is the amount from one wing. Underlined are the individuals that showed indels in the targeted genomic regions based on DNA extracted from the thorax. Numbers are the individual identification, L and R are left and right wings respectively.

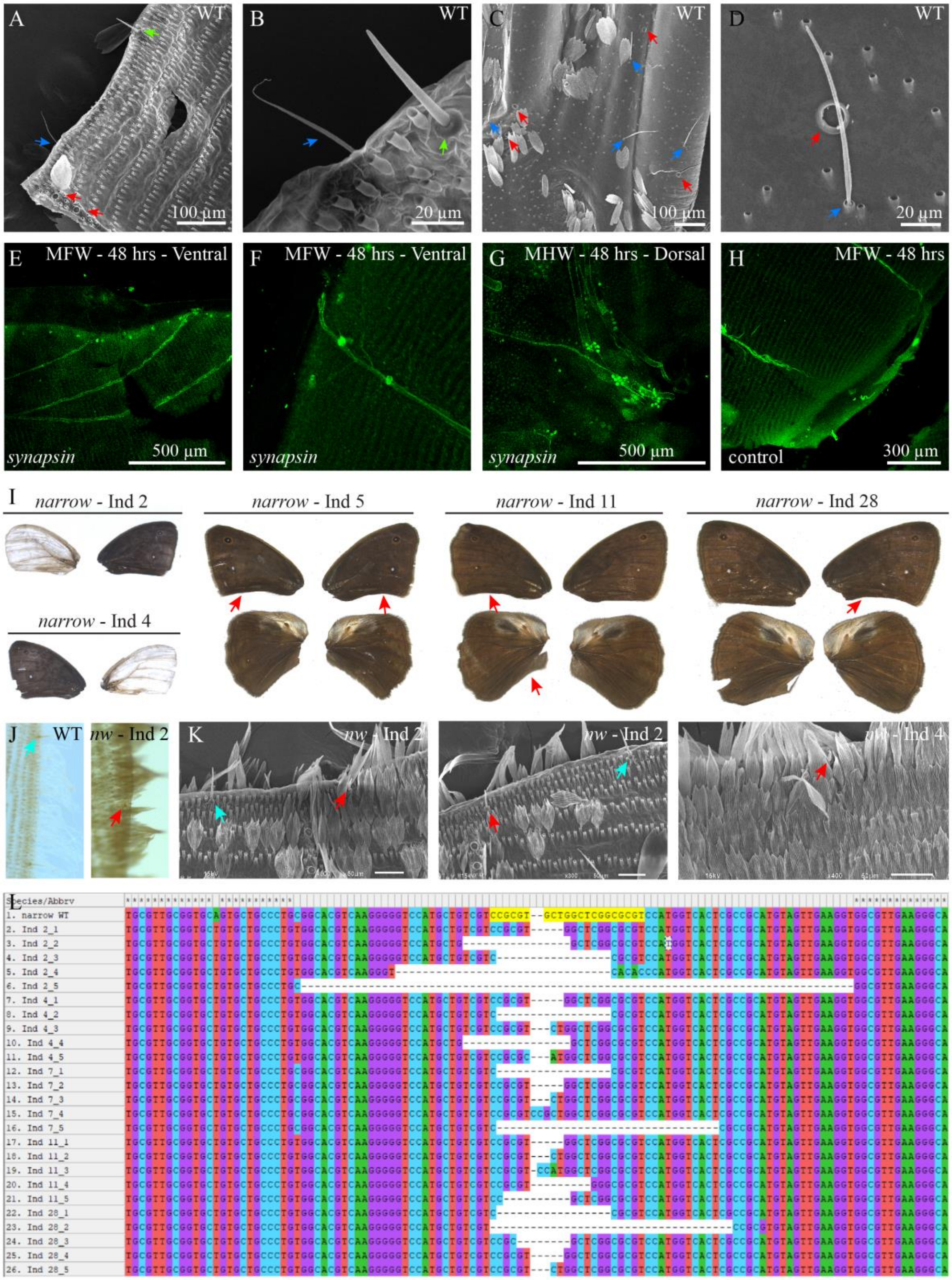


**Figure S16: Sensory cell types on *B. anynana* wings and *narrow* crispant phenotypes.** (A-D) Different sensory cell types on *B. anynana* wings. (A) At the margin, stout mechanosensory bristles are located on the ventral surface (green arrow) while thin, hair-like bristles are located on the dorsal surface (blue arrow). Campaniform sensilla are seen in a characteristic pattern on the marginal ends of the ventral veins (red arrows). (B) Magnified view of the ventral (green arrow) and dorsal (blue arrow) marginal sensory cell types. (C) Sensory cell types at the base of the wing near the hinge region. Red arrows indicate campaniform sensilla while the blue arrows indicate hair-like sensory bristles. (D) Magnified view of a campaniform sensilla (red arrow) and the hair-like sensory bristle (blue arrow). (E-G) Synapsin protein expression is seen in the innervated sensory cell types (E) along the anterior margin, (F) corresponding to the two campaniform sensilla on the ventral veins near the margin and (G) at the base of the wing near the hinge region. (H) Control stains show no synapsin expression in these cells. (I) Representative images of the phenotypes obtained upon *narrow* knockout in *B. anynana*. Red arrows indicate changes in wing shape. (J, K) *narrow* crispant individuals exhibit smaller marginal mechanosensory bristles (red arrows) as compared to the wildtype/ wildtype-like bristles in *narrow* crispant wings (blue arrows). Scale bars in K are 50 µm. (L) Indels at the targeted locus in various crispant individuals. MFW: Male forewing; MHW: Male hindwing.

**Table S1**: Top usage genes annotated_k13_0.18threshold. Found in a separate excel file.

**Table S2**: Histone and ribosomal proteins removed from the gene count matrix

| Type | Genes (NCBI Gene IDs) |
|---|---|
| Histones | "112054639", "112049474", "112057469", "112051713", "112049472", "112051725", "112049513", "112051712", "112049497", "112049516", "112051714", "112050938", "112050939", "112057467", "112049496", "112049476", "112056552", "112051726" |
| Ribosomal proteins | "112045345", "112042909", "112044081", "112055175", "112047303", "112048726", "112052451", "112052453", "112049145", "112048475", "112047442", "112046298", "112055063", "112051774", "112053219", "112057881", "112051107", "112051307", "112044661", "112046041", "112048941", "112051258", "112043474", "112044824", "112052195", "112043308", "112051236", "112058575", "112048815", "112049701", "112055280", "112049204", "112052391", "112049437", "112046504", "112046748", "112048857", "112052026", "112054272", "112050736", "112053859", "112044884", "112046892", "112055487", "112044484", "112045583", "112052997", "112056026", "112053495", "112048248", "112049427", "112052210", "112056661" |

**Table S3**: List of primers (*in situ* hybridization)

| Gene (NCBI Gene ID) | Primer Name | Primer Sequence | |
|---|---|---|---|
| *cpo* (112046215) | AM 1979 | Forward | 5' GGACTCCGTCAACACCGC 3' |
| | AM 1980 | Reverse | 5' GATGGCGGGATGGACCAG 3' |
| *HR38* (112044883) | AM 1379 | Forward | 5' GGAGATACGGCTGCATGTC 3' |
| | AM 1380 | Reverse | 5' CTAGGACTCGGCTGAAGTG 3' |
| *senseless* (112050060) | AM 1383 | Forward | 5' GTGAAGAGATGCCGCAGC 3' |
| | AM 1384 | Reverse | 5' GCTTGTTCCTCGCCGTAG 3' |
| *rst-scale cells* (112046704) | AM1983 | Forward | 5' GTTTGGAACGCTATGTGCTG 3' |
| | AM1984 | Reverse | 5' CATATAACAGAGAGCGAGGC 3' |

**Table S4**: List of guide RNA sequences for CRISPR-Cas9

| Gene (NCBI Gene ID) | Primer Name | Primer Sequence |
|---|---|---|
| *HR38* (112044883) | AM1457 | GAAATTAATACGACTCACTATAGGAATGGTGAAAGAGGTCGTG GTTTTAGAGCTAGAAATAGC |
| *senseless* (112050060) | AM1538 | GAAATTAATACGACTCACTATAGGAGTGGATGAGCAGATGCG GTTTTAGAGCTAGAAATAGC |
| *Pax5* (112056518) | AM1399 | GAAATTAATACGACTCACTATAGGTAGCGACCCCTCCTGTGG GTTTTAGAGCTAGAAATAGC |
| *paramyosin* (112046058) | AM1542 | GAAATTAATACGACTCACTATAGGTAAGGCTGAAGTCCAAAGGT GTTTTAGAGCTAGAAATAGC |
| *narrow* (112055826) | AM1544 | GAAATTAATACGACTCACTATAGGACGCGCCGAGCCAGCACG GTTTTAGAGCTAGAAATAGC |
| *circadian clock* (112043107) | AM1690 | GAAATTAATACGACTCACTATAGGTTTGCCAGAATGGGGAGTGG GTTTTAGAGCTAGAAATAGC |
| *grainyhead* (112046894) | AM1347 | GAAATTAATACGACTCACTATAGGCGAGCAGCAGTTCCCAGAGG GTTTTAGAGCTAGAAATAGC |
| *FAR* (112046326) | AM1998 | GAAATTAATACGACTCACTATAGGTAGGAGCGTCCTAGTGAC GTTTTAGAGCTAGAAATAGC |

**Table S5**: Genotyping primers for crispants

| Gene (NCBI Gene ID) | Primer Name | Primer Sequence (Reading primer + gene specific primer) |
|---|---|---|
| *HR38* (112044883) | AM 1868<br>AM 1869 | Forward<br>ACACTCTTTCCCTACACGACGCTCTTCCGATCTCGTGTCCAGTGGATAAGAG<br>Reverse<br>GTGACTGGAGTTCAGACGTGTGCTCTTCCGATCTGTTCGGCTCTCTGTATTGTG |
| *senseless* (112050060) | AM 1964<br>AM 1965 | Forward<br>ACACTCTTTCCCTACACGACGCTCTTCCGATCTCCAATGACAAGCATAGTG<br>Reverse<br>GTGACTGGAGTTCAGACGTGTGCTCTTCCGATCTGATGATGATGACAAATACC |
| *Pax5* (112056518) | AM 1968<br>AM 1969 | Forward<br>ACACTCTTTCCCTACACGACGCTCTTCCGATCTCTAGCCCGAAGTTAGAAAC<br>Reverse<br>GTGACTGGAGTTCAGACGTGTGCTCTTCCGATCTGATACTACGAAACGGGGAG |
| *paramyosin* (112046058) | AM 1870<br>AM 1871 | Forward<br>ACACTCTTTCCCTACACGACGCTCTTCCGATCTCATTTCAGGAACAAAGATTACAC<br>Reverse<br>GTGACTGGAGTTCAGACGTGTGCTCTTCCGATCTCTAACTTCTCACATTGTATTTCC |

| | | |
|---|---|---|
| *narrow* (112055826) | AM 1872<br>AM 1873 | Forward<br>ACACTCTTTCCCTACACGACGCTCTTCCGATCTCAACAAGAGGCCAATGGAG<br>Reverse<br>GTGACTGGAGTTCAGACGTGTGCTCTTCCGATCTGAATTTTCTCATCGCAGGATAC |
| *circadian clock* (112043107) | AM 1938<br>AM 1939 | Forward<br>ACACTCTTTCCCTACACGACGCTCTTCCGATCTCACCTTGATAAGTGTAGACTG<br>Reverse<br>GTGACTGGAGTTCAGACGTGTGCTCTTCCGATCTCCCGGCTGGAACTTTTTTAG |
| *grainyhead* (112046894) | AM 1940<br>AM 1941 | Forward<br>ACACTCTTTCCCTACACGACGCTCTTCCGATCTGTGCAGATTTTCACCATACGG<br>Reverse<br>GTGACTGGAGTTCAGACGTGTGCTCTTCCGATCTGTATAGTGTACTGGCAGACG |
| *FAR* (112046326) | AM 1999<br>AM 2000 | Forward<br>ACACTCTTTCCCTACACGACGCTCTTCCGATCTCACCGACTTCACAATTGTG<br>Reverse<br>GTGACTGGAGTTCAGACGTGTGCTCTTCCGATCTGTTGCATGTCAGCGAACC |
